# Lichen symbiosis does not impose a uniform genomic syndrome on algae

**DOI:** 10.64898/2026.07.23.740355

**Authors:** Abigail R. Meyer

## Abstract

Symbiosis has lasting effects on the genomic evolution of interacting organisms. Shifting niches and selection intensities are well-documented drivers of adaptive evolution, but nonadaptive evolution is an equally important, if less conspicuous, dimension of symbiont genome evolution. Through a reduction in effective population size (Ne), symbiosis can reduce the efficacy of natural selection in symbionts; however, evidence of this “genomic syndrome” is derived almost entirely from bacteria. Whether it extends to eukaryotic symbioses with complex demography remains unclear. Here, lichen-forming algae are examined for signatures of a genome-wide reduction in the efficacy of selection arising from demographic shifts associated with symbiosis. To test this, four lichen-forming algal taxa were compared to their closest free-living relatives using complementary measures of (i) the strength of molecular evolution (dN/dS, K), (ii) codon use bias (ENC’), and (iii) base composition at synonymous sites (GC3). Lichen-forming algae showed heterogeneous responses across all signatures of molecular evolution examined. The effect of lifestyle (lichen-forming vs. free-living) on dN/dS ratios and codon use bias was lineage specific. GC3 was uniformly reduced in lichen-forming taxa, but the underlying causes differed, and no genes showed uniformly intensified or relaxed selection across all four lichen-forming taxa. These results suggest that, while symbiosis often reshapes symbiont population genetics in ways that elevate the role of drift, lichen-forming algae do not show a uniform reduction in the efficacy of selection. Rather, the demographic effects and functional demands of each specific lichen symbiosis likely shape genome evolution in lineage-specific ways.

## Introduction

Symbiosis is a widely recognized driver of genomic evolution for both hosts and their microbial symbionts (Husnik & McCutcheon, 2018; McCutcheon & Moran, 2012; Moran, 2007; Wernegreen, 2002). These persistent, interspecies interactions can introduce genes (e.g. horizontal gene transfer, Keeling & Palmer, 2008) combine lineages (e.g. eukaryogenesis, Sagan, 1967), accelerate or slow the pace of molecular evolution (Moran, 1996; Woolfit & Bromham, 2003) relax selective constraints (Douglas, 2016), and drive adaptive divergence (Sudakaran et al., 2017). Why symbiont genomes evolve faster, or slower, than those of their free-living relatives remains an open question in evolutionary biology and is often attributed to the interplay of ecological dependence, coevolution, and demographic shifts (Bromham, 2009).

Symbiosis alters the population genetic environment of symbionts, in part, through reductions in effective population size (Ne). Reduced Ne decreases the efficacy of natural selection and strengthens the role of genetic drift in influencing allele frequencies across generations, increasing the probability of mildly deleterious mutations drifting to fixation (Lynch, 2007b; Lynch & Conery, 2003; Ohta, 1992). Across bacterial symbionts, signatures of reduced efficacy of selection are diagnostic features of the pervasiveness of Ne reduction, ranging from extreme and sustained in obligate, vertically transmitted endosymbionts (Bennett & Moran, 2015; Moran, 1996; Moran et al., 2008; Wernegreen, 2002) to weaker in systems with mixed transmission modes, ongoing recombination, or retained free-living phases (Russell et al., 2017, 2020).

Bacterial symbionts have thus become a model for how demographic constraint interacts with the efficacy of selection; however, far fewer studies have investigated whether similar demographic constraints affect genome-wide evolution in eukaryotic symbionts. Early work using single-locus sequences in lichen-forming and endosymbiotic fungi showed elevated rates of molecular evolution (Lutzoni & Pagel, 1997; Woolfit & Bromham, 2003). In genome-scale studies, some systems show elevated substitution rates and reduced efficacy of selection consistent with reduced Ne (Conlon et al., 2021; Haag et al., 2020), whereas others, including lichen-forming fungi, show slower evolutionary rates (Ametrano et al., 2022). Continuing to characterize molecular evolutionary patterns across eukaryotic symbioses is necessary to determine whether the signatures of Ne reduction characterized in bacterial symbiont genomes are transferable to systems with fundamentally different modes of reproduction, mutational dynamics, and genome architectures (Lynch, 2007b).

Lichen symbioses offer an opportunity to assess whether demographic constraints on effective population size have shaped genome evolution in an extracellular eukaryotic symbiosis. Lichens symbioses are partnerships composed of a lichen-forming fungus, a lichen-forming “alga” (green algae or cyanobacteria), and a myriad of additional microorganisms (secondary fungi and algae, bacteria, microinvertebrates, etc.). Lichen symbioses are extracellular and span a continuum of transmission modes, partner fidelity (e.g. specificity/selectivity) and dependence on symbiosis. While historically studied from the fungal perspective, recent research has brought increased attention to the ecology (Koch et al., 2023), diversity (Sanders & Masumoto, 2021; Veselá et al., 2024) and evolution (Nelsen et al., 2021, 2022; Puginier et al., 2024) of lichen-forming algae. Comparative genomics has revealed gene family expansions (e.g. glucose/ribitol dehydrogenase and short chain dehydrogenase reductase) and candidate symbiosis associated genes (e.g. glycoside hydrolase 8), potentially underpinning the capacity for algae to transfer carbon and mediate fungal cell wall remodeling at the symbiotic interface (Armaleo et al., 2019; Poquita-Du et al., 2024; Puginier et al., 2024; Xiong et al., 2025).

Despite recent advances in understanding the ecology and functional genomic evolution of lichen-forming algae, how symbiosis affects their demography remains poorly understood. Several features of lichen-forming algal biology suggest that Ne may be reduced through symbiosis. Because algae live entangled within or beneath fungal tissues, gene flow may be restricted among algae, forming spatially or genetically isolated subpopulations. Their reproductive biology may also impose recurrent bottlenecks, resulting in a winnowing of genetic diversity at each generation (Fig. 1). Such demographic fluctuations, with periods of low Ne during transmission followed by expansion in or outside of symbiosis, can have lasting genomic effects because even brief reductions in Ne disproportionately reduce long-term Ne (Wright 1938). If lichen symbiosis reduces algal Ne through demographic bottlenecks, population genetic theory predicts genome-wide reduction in the efficacy of selection (Ohta, 1992).

**Figure 1.**
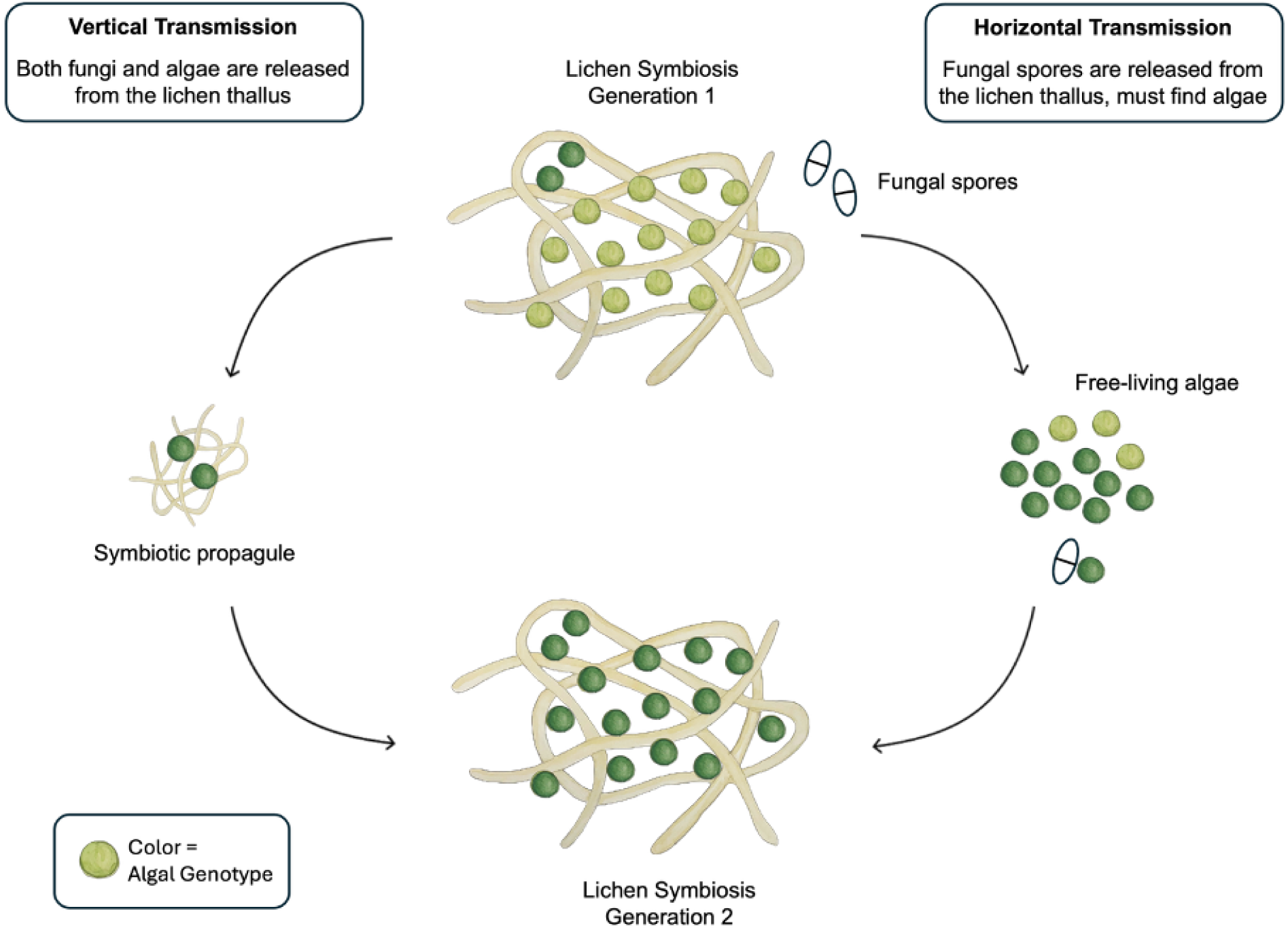
Conceptual model illustrating the proposed population bottlenecks that occur during lichen transmission and resynthesis across generations. In vertical transmission, dispersal of a small subset of algal cells can influence population genetic diversity of future lichen generations. In horizontal transmission, the algal cells, either free-living or lichenized, that are acquired by fungal spores during resynthesis represent a founder population, similarly shaping the genetic diversity of future symbioses.

**Alt text:** Lifecycle diagram demonstrating that in both vertical and horizontal transmission each new lichen generation inherits only a subset of the algal genotypes present in the previous generation.

To investigate how lichen symbiosis affects the molecular evolution of algae, a comparative genomics approach was used to examine genomic signatures across four pairs of lichen-forming and closely related free-living algae. These signatures include four complementary analyses to detect selection: dN/dS ratios to assess overall strength of purifying selection, K parameters (RELAX) to test for relaxation or intensification of selection, ENC’ (effective number of codons corrected for GC content) as an indicator of codon use bias (CUB), and GC content at the third codon position (GC3) to assess compositional shifts at synonymous sites. If lichen symbiosis reduces Ne, it is predicted that lichen-forming algae will exhibit higher dN/dS (reduced efficacy of selection at nonsynonymous sites; Ohta 1992), K < 1 (relaxed selection; Wertheim et al. 2015), higher ENC’ (reduced CUB; Bulmer 1991; Sharp et al. 2010), and lower GC3 without an associated GC12 shift (reduced selection at synonymous sites rather than shift in mutational spectrum; Sueoka 1988). If lichen symbiosis does not cause a uniform reduction in Ne and resultant reduction in the efficacy of selection, heterogeneity in these molecular signatures across taxa is expected, potentially reflecting lineage-specific differences in bottleneck severity and the unique functional demands of each lichen symbiosis.

## Results

To investigate differences in rates of molecular evolution between lichen-forming and free-living algae, seven publicly available genomes were analyzed (Fig. 2A, Table 1). Multiple representative genomes are available for both *Trebouxia* and *Asterochloris*, and the results were shown to be robust to genome choice (Supplementary Fig 1). *Trebouxia* and *Asterochloris* share the same closest free-living relative, making the *Trebouxia* vs. *Myrmecia* and *Asterochloris* vs*. Myrmecia* comparisons not fully independent.

**Figure 2.**
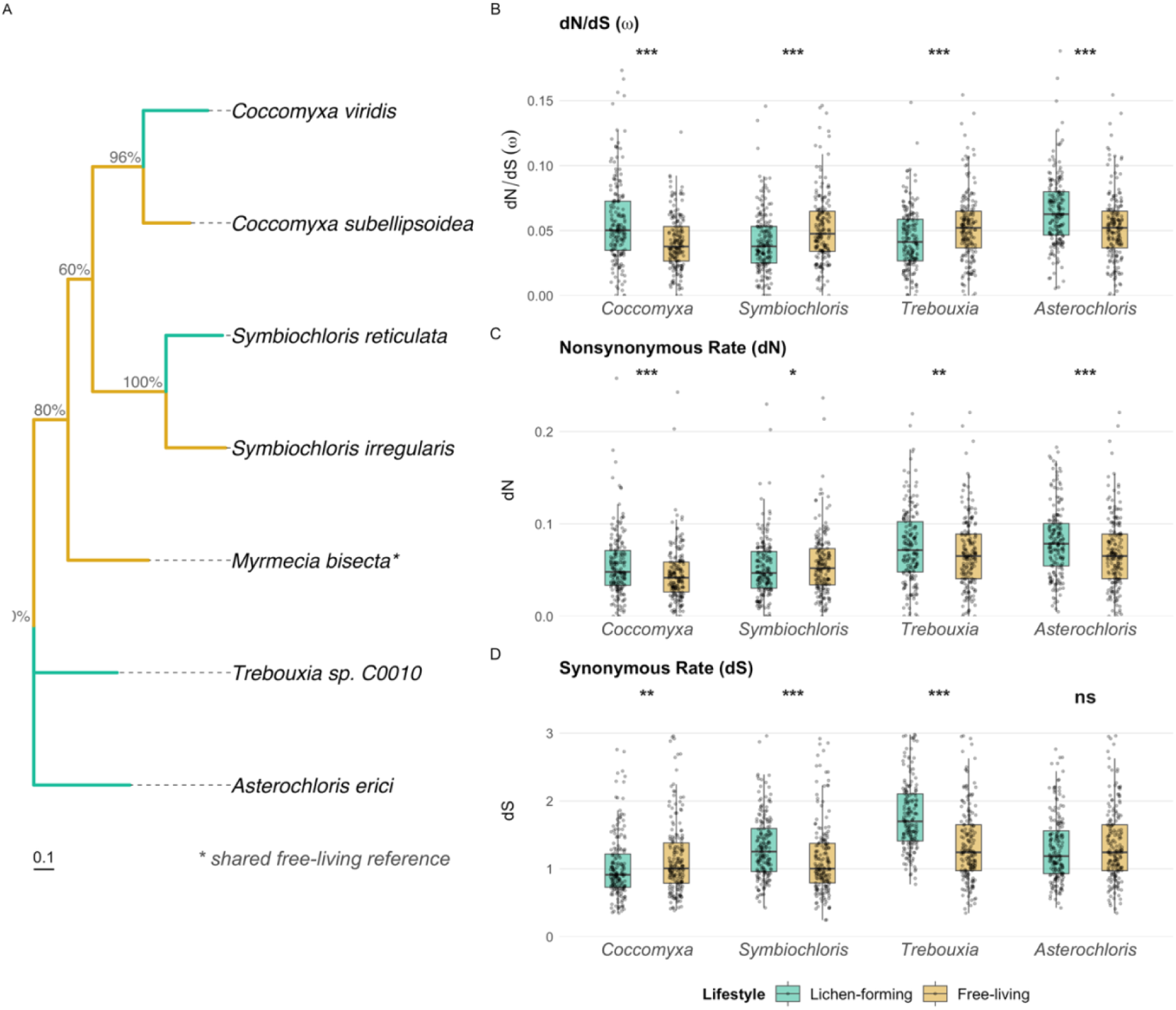
Rates of molecular evolution in lichen-forming and free-living algae. (A) Phylogenetic relationship (unrooted consensus species tree based on 951 gene trees) of the algal taxa examined in this study. Node values represent the percentage of gene trees that support each split. Scale bar represents 0.1 substitutions/site. (B) Relative dN/dS ratio (ω) (C) Absolute nonsynonymous substitution rates (dN) (D) Absolute synonymous substitution rates (dS) across taxon pairs. Points represent individual SOGs. Significance levels from pairwise contrasts of estimated marginal means are indicated (*p < 0.05, **p < 0.01, ***p < 0.001).

**Table 1.**
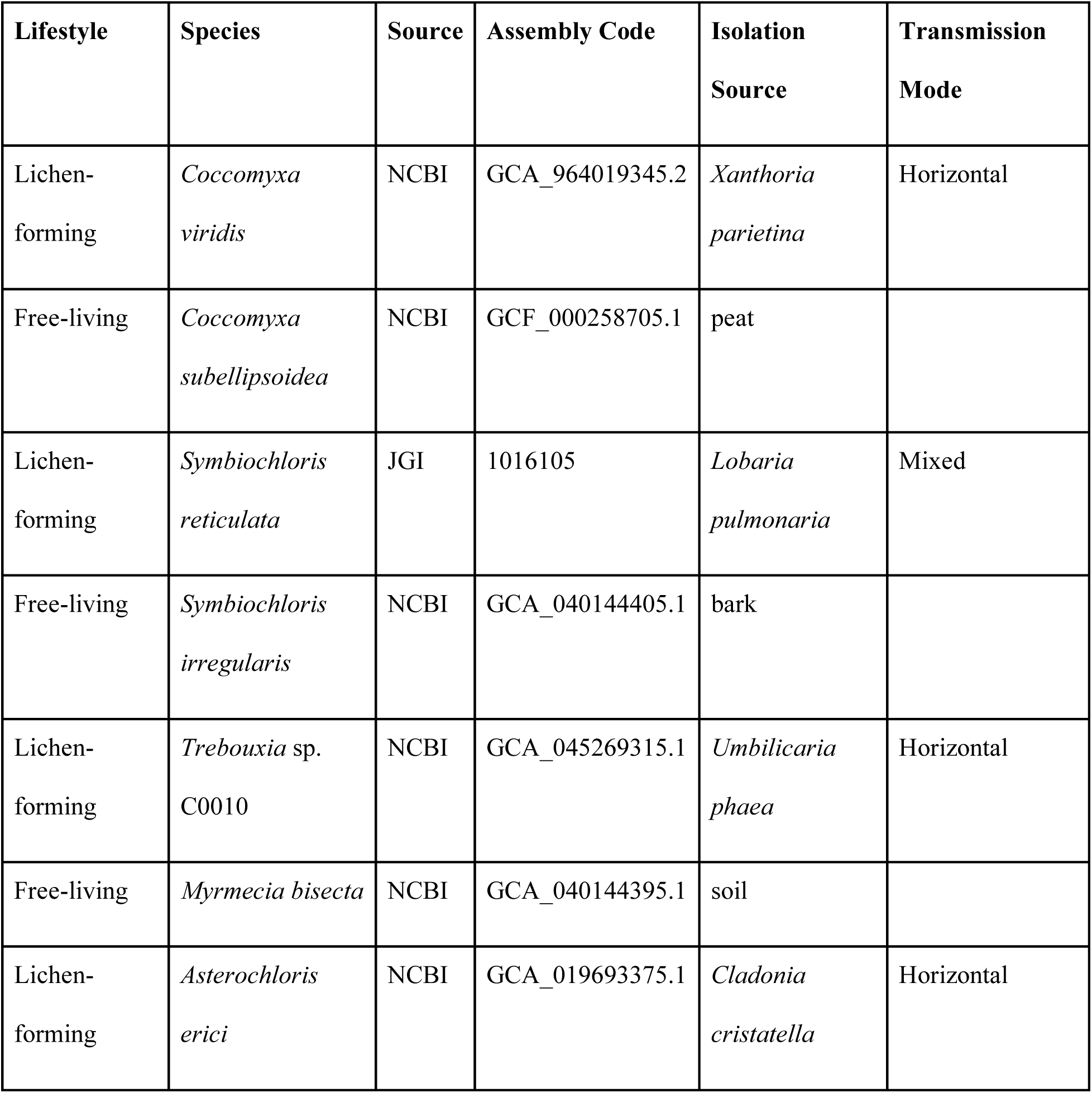
Table of taxa used in this study, including their lifestyle, specific genomic resource used, isolation source and transmission mode. Lifestyle designations reflect each taxon’s predominant life-history and the provenance of the specific genomic resource used. *Cocccomyxa subellipsoidea* is associated with the lichen-forming fungus *Lichenomphalia spp.* (Zoller & Lutzoni, 2003); however, *C. subellipsoidea* is overwhelmingly free-living, and the specific genome used here is from a free-living collection. Transmission mode is based on the taxonomic characteristics of the specific lichen symbiosis (i.e., lichen-forming fungi) from which the algae was isolated. Mechanisms of algal acquisition remain poorly understood; however, because thallus fragmentation is thought to be an important transmission vector (Sanders, 2023), most associations likely involve some mixture of vertical and horizontal transmission.

| Lifestyle | Species | Source | Assembly Code | Isolation Source | Transmission Mode |
| --- | --- | --- | --- | --- | --- |
| Lichen-forming | *Coccomyxa viridis* | NCBI | GCA_964019345.2 | *Xanthoria parietina* | Horizontal |
| Free-living | *Coccomyxa subellipsoidea* | NCBI | GCF_000258705.1 | peat |  |
| Lichen-forming | *Symbiochloris reticulata* | JGI | 1016105 | *Lobaria pulmonaria* | Mixed |
| Free-living | *Symbiochloris irregularis* | NCBI | GCA_040144405.1 | bark |  |
| Lichen-forming | *Trebouxia* sp.<br>C0010 | NCBI | GCA_045269315.1 | *Umbilicaria phaea* | Horizontal |
| Free-living | *Myrmecia bisecta* | NCBI | GCA_040144395.1 | soil |  |
| Lichen-forming | *Asterochloris erici* | NCBI | GCA_019693375.1 | *Cladonia cristatella* | Horizontal |

### Rates of Molecular Evolution

To compare rates of molecular evolution between species, 951 single copy orthologous genes (SOGs) shared across all seven taxa were identified. Codon based alignments and gene trees for each SOG were used to estimate the ratio of nonsynonymous to synonymous substitutions between species (dN/dS). This parameter was estimated via maximum likelihood using a model that fit a separate dN/dS ratio (ω) for each branch (codeml “free-ratio” model; Yang, 2007). After filtering for extreme or unreliable estimates (0.01 < dS < 3, dN/dS < 10; see Methods), 185 SOGs remained for analysis. Estimated lifestyle effects were robust to these filtering thresholds (Supplementary Fig. 2).

A linear mixed-effects model was fit to test whether lifestyle (lichen-forming vs. free-living) affected ω. The effect of lifestyle on ω differed significantly across taxon pairs, with a significant lifestyle × taxon pair interaction (F_3,1288_ = 46.43, p < 0.0001; Fig. 2B, Table 2). Pairwise contrasts of estimated marginal means showed two pairs with significantly elevated ω in lichen-forming species (*Coccomyxa*: Δω = +0.0155; *Asterochloris*: Δω = +0.0125; both p < 0.0001) and two pairs with significantly reduced ω in lichen-forming species (*Symbiochloris*: Δω = −0.0114; *Trebouxia*: Δω = −0.0090; both p < 0.0001) relative to their free-living counterparts. The estimated marginal means averaged across taxon pairs (pooled contrast from the lifestyle × taxon-pair interaction model) showed a small, non-significant difference between lifestyles (Δω = +0.0019, p =0.0681), reflecting opposing directional effects across taxa comparisons that cancel when pooled.

**Table 2.** Table of estimated marginal mean differences in dN, dS, and ω between lichen-forming and free-living algae within each comparison, derived from separate linear mixed-effects models with fixed lifestyle × taxon pair interaction and SOG as a random intercept. P-values are from pairwise contrasts of estimated marginal means and are Bonferroni-adjusted across the four pairs.

| Lichen-forming<br>algae | Free-living<br>algae | $\Delta dN$<br>(SE, p-value) | $\Delta dS$<br>(SE, p-value) | $\Delta \omega$<br>(SE, p-value) |
| --- | --- | --- | --- | --- |
| *Coccomyxa*<br>*viridis* | *Coccomyxa*<br>*subellipsoidea* | +0.0102<br>(0.0021, <0.0001) | -0.1184<br>(0.0413, 0.0042) | +0.0155<br>(0.0021, <0.0001) |
| *Symbiochloris*<br>*reticulata* | *Symbiochloris*<br>*irregularis* | -0.0048<br>(0.0021, 0.0226) | +0.1759<br>(0.0413, <0.0001) | -0.0114<br>(0.0021, <0.0001) |
| *Trebouxia* sp.<br>C0010 | *Myrmecia*<br>*bisecta* | +0.0064<br>(0.0021, 0.0021) | +0.4311<br>(0.0413, <0.0001) | -0.0090<br>(0.0021, <0.0001) |
| *Asterochloris*<br>*erici* | *Myrmecia*<br>*bisecta* | +0.0134<br>(0.0021, <0.0001) | -0.0472<br>(0.0413, 0.2533) | +0.0125<br>(0.0021, <0.0001) |

To understand these differences across taxon pairs, absolute dN and dS values were also examined. Nonsynonymous substitution rates (dN) were significantly elevated in lichen-forming lineages for three of four taxon pairs (*Asterochloris*: ΔdN = +0.01342, p < 0.0001; *Coccomyxa*: ΔdN = +0.01015, p < 0.0001; *Trebouxia*: ΔdN = +0.00641, p = 0.0021), while *Symbiochloris* showed lower dN (ΔdN = −0.00475, p = 0.0226) (Fig. 2C, Table 2). Synonymous substitution rates (dS) between taxon pairs showed greater heterogeneity across free-living and lichen-forming comparisons: *Symbiochloris* and *Trebouxia* exhibited significant increases (*Symbiochloris*: ΔdS = +0.1759, p < 0.0001; *Trebouxia*: ΔdS = +0.4311, p < 0.0001), *Coccomyxa* showed a significant decrease (ΔdS = −0.1184, p = 0.0042) and *Asterochloris* showed no significant difference (ΔdS = −0.0472, p = 0.2533) (Fig. 2D, Table 2).

To assess whether ΔdS heterogeneity could be attributed to differences in evolutionary divergence time, phylogenetic distances between species pairs were examined. Phylogenetic distances ranged from 0.54-0.57 substitutions/site for within-genus comparisons (*Coccomyxa* and *Symbiochloris*) to 0.97-1.03 substitutions/site for cross-genus comparisons (*Trebouxia* and *Asterochloris* vs. *Myrmecia*). The two most divergent pairs showed opposite dS patterns (*Trebouxia*: ΔdS = +0.43; *Asterochloris*: ΔdS = −0.05; Supplementary Fig. 3).

**Alt text:** Species tree of taxa in the study and their absolute and relative rates of molecular evolution. Two taxon pairs show higher ratios of nonsynonymous to synonymous substitutions in the lichen-forming taxon and two show lower. Nonsynonymous and synonymous rates shift in inconsistent directions across pairs.

### Codon Use Bias

To quantify CUB while accounting for differences in background nucleotide composition between taxa, the effective number of codons corrected for GC content (ENC’; Novembre, 2002) was examined. The effect of lifestyle on ENC’ varied significantly across pairs (lifestyle × taxon pair interaction: F^3,6650^ = 438.09, p < 0.0001; Fig. 3A). Relative to their free-living counterparts, *Trebouxia* showed the largest increase in ENC’ (ΔENC’ = +2.9568, p < 0.001) and *Symbiochloris* showed a small increase (ΔENC’ = +0.9400, p < 0.001). In contrast, *Coccomyxa* showed a significant but small decrease in ENC’ (ΔENC’ = −0.5974, p < 0.001), and *Asterochloris* showed no significant difference (ΔENC’ = −0.0712, p = 0.3421).

**Figure 3.**
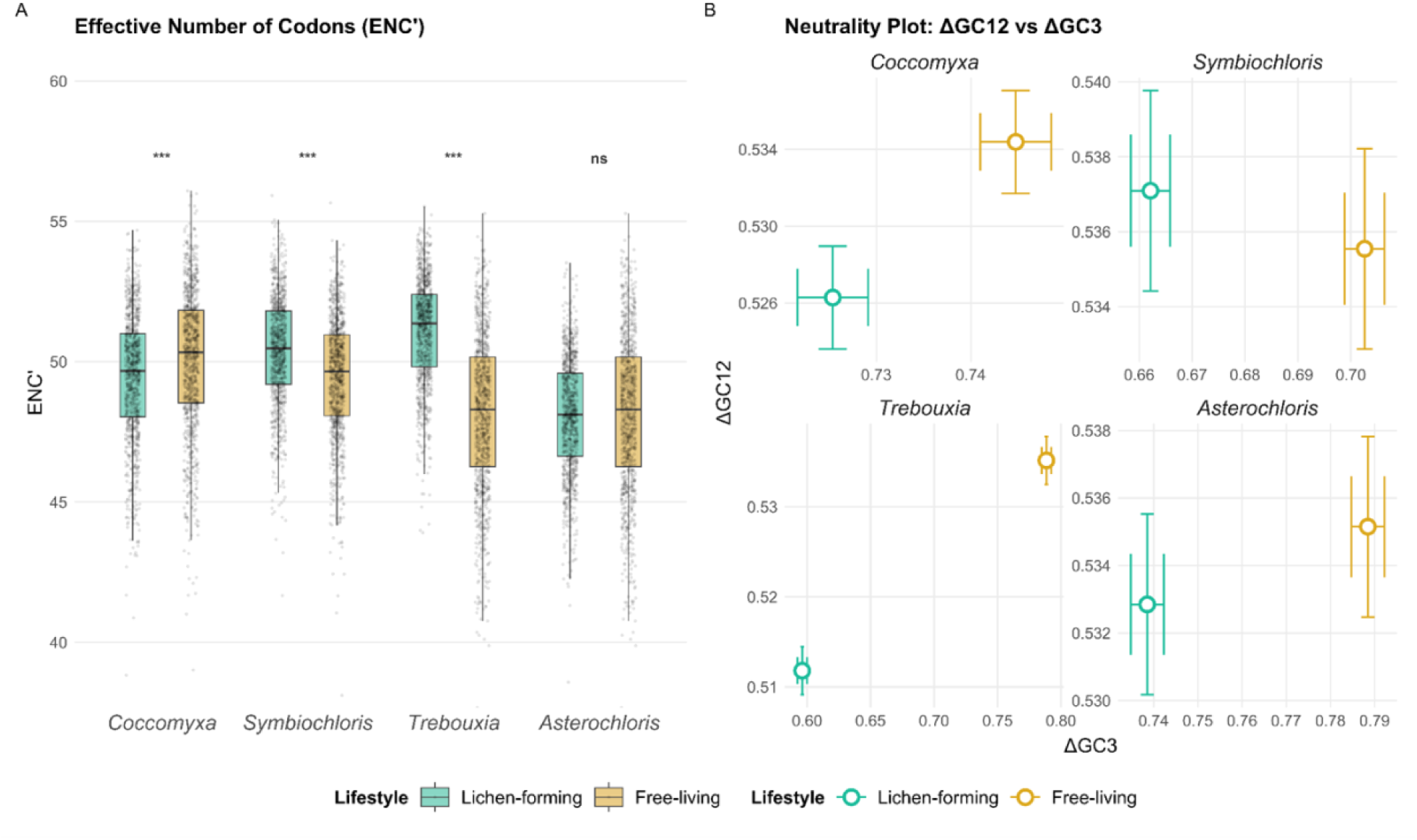
Codon use bias (CUB) in lichen-forming and free-living algae. (A) Effective number of codons (ENC’) (B) Neutrality plot of estimated marginal mean GC content at first and second codon positions (GC12) against estimated marginal mean GC content at the third codon position (GC3), shown as centroids (± CI) for each species within each taxon pair. Displacement between lichen-forming and free-living centroids that is predominantly horizontal (shift in GC3 with similar GC12) is consistent with reduced efficacy of selection on synonymous codon usage rather than a genome-wide shift in mutational pressure. A vertical shift in centroid displacement suggests differences in genome-wide mutational pressure contributes to GC3 differences between taxon pairs. Significance levels from pairwise contrasts of estimated marginal means are indicated (*p < 0.05, **p < 0.01, ***p < 0.001).

To assess whether dS variation reflects changes in CUB, the relationship between ΔdS and ΔENC’ across taxon pairs was examined. ΔENC’ and ΔdS covaried in sign and rank order across all four comparisons. Both *Trebouxia* (ΔENC’ = +2.9568, ΔdS = +0.4311) and *Symbiochloris* (ΔENC’ = +0.9400, ΔdS = +0.1759) had significantly elevated ENC’ and dS, whereas *Coccomyxa* had a lower ENC’ and dS (ΔENC’ = −0.5974, ΔdS = −0.1184). Neither ENC’ nor dS were significantly different in *Asterochloris* (ΔENC’ = −0.0712, ΔdS = −0.0472) (Table 3). An n of 4 for these comparisons limits formal correlation testing between ΔdS and ΔENC’.

**Table 3.** Table of estimated marginal mean differences in ENC’, GC12, and GC3 between lichen-forming and free-living algae within each comparison, derived from separate linear mixed-effects models with fixed lifestyle × taxon pair interaction and SOG as a random intercept. P-values are from pairwise contrasts of estimated marginal means and are Bonferroni-adjusted across the four pairs.

| Lichen-forming<br>algae | Free-living<br>algae | ΔENC'<br>(SE, p-value) | ΔGC12<br>(SE, p-value) | ΔGC3<br>(SE, p-value) |
| --- | --- | --- | --- | --- |
| *Coccomyxa*<br>*viridis* | *Coccomyxa*<br>*subellipsoidea* | -0.5974<br>(0.0750, <0.0001) | -0.0081<br>(0.0008, <0.0001) | -0.0194<br>(0.0019, <0.0001) |
| *Symbiochloris*<br>*reticulata* | *Symbiochloris*<br>*irregularis* | +0.9400<br>(0.0750, <0.0001) | +0.0016<br>(0.0008, 0.0626) | -0.0405<br>(0.0019, <0.0001) |
| *Trebouxia* sp.<br>C0010 | *Myrmecia*<br>*bisecta* | +2.9568<br>(0.0750, <0.0001) | -0.0233<br>(0.0008, <0.0001) | -0.1924<br>(0.0019, <0.0001) |
| *Asterochloris*<br>*erici* | *Myrmecia*<br>*bisecta* | -0.0712<br>(0.0750, 1.0000) | -0.0023<br>(0.0008, 0.0057) | -0.0499<br>(0.0019, <0.0001) |

GC3 showed a uniform reduction in lichen-forming taxa across all four comparisons (all p < 0.0001; Fig. 3, Table 3). To assess if shifts in GC3 reflect genome-wide changes in the mutational spectrum or shifts in the efficacy of selection at synonymous sites, the relationship between the mean GC content at the first and second codon position (GC12) and GC3 within each taxon pair was examined (Fig. 3B). If directional mutational pressure drives GC3 shifts, GC12 and GC3 should shift proportionately (Sueoka, 1988). GC12 was significantly lower in lichen-forming *Coccomyxa* (ΔGC12 = −0.0081, p < 0.001) indicating a genome-wide compositional shift in GC content contributes to lower GC3. In *Trebouxia* GC12 decreased relative to its free-living counterpart, however the magnitude of the GC3 shift (ΔGC3 = −0.1924) was substantially larger than the GC12 shift (ΔGC12 = −0.0233). In *Symbiochloris*, GC12 did not differ between lifestyles (ΔGC12 = +0.0016, p = 0.063), consistent with the GC3 decrease being driven by changes at synonymous sites (i.e. reduced efficacy of selection on CUB). *Asterochloris* showed a small but significant decrease in GC12 (ΔGC12 = −0.0023, p = 0.006) relative to its free-living counterpart.

**Alt text:** Box plots of effective number of codons and a neutrality plot showing relative shift in average GC content at the first and second codon position versus the third codon position. Lichen-forming *Trebouxia* and *Symbiochloris* shift toward less biased codon usage. All lichen-forming centroids move toward lower GC content at the third codon position. *Symbiochloris*, *Asterochloris*, and *Trebouxia* shift mainly at the third codon position, while *Coccomyxa* also shows a substantial shift at the first and second position.

### Selection Intensity (RELAX)

To test for changes in selection intensity between lichen-forming and free-living algae, RELAX was applied separately to each SOG for each of four taxon-pair comparisons. In each comparison, the lichen-forming taxon was labeled as the test branch and its free-living counterpart as the reference branch, with all other taxa unlabeled. RELAX failed to converge for a subset of SOG analyses, leading to variability in gene number across taxa (Table 4). RELAX estimates a relaxation parameter (K), where K < 1 indicates relaxation and K > 1 indicates intensification of selection. The majority of genes in lichen-forming taxa showed no significant change in selection intensity. Of those that were significant (p < 0.05), *Trebouxia* sp. C0010 and *Symbiochloris reticulata* showed a significant excess of genes under intensification of selection (binomial test, p < 0.0001 and p = 0.031, respectively). *Coccomyxa viridis* and *Asterochloris erici* did not show a significant bias towards intensification or relaxation of selection (p = 0.254 and p = 1.000, respectively) (Fig. 4, Table 4).

**Figure 4.**
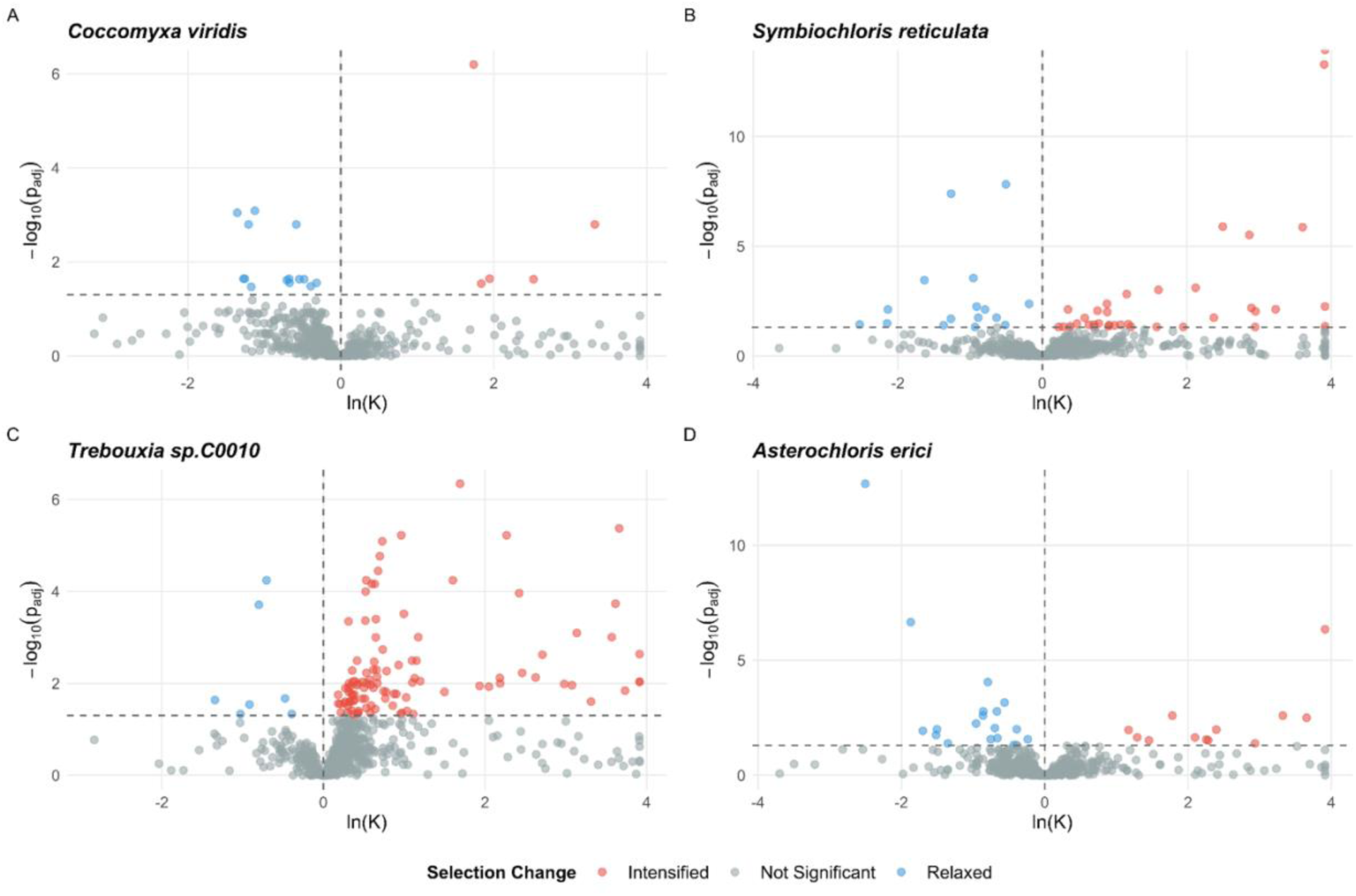
Shifts in selection intensity in lichen-forming algae estimated using RELAX. Volcano plots show -log^10^ of the Benjamini–Hochberg adjusted p-value against the natural log of the selection intensity parameter (K) for (A) *Coccomyxa viridis*, (B) *Symbiochloris reticulata*, (C) *Trebouxia* sp. C0010, and (D) *Asterochloris erici*. ln(*K*) > 0 indicates intensified selection and ln(*K*) < 0 indicates relaxed selection in each lichen-forming taxon relative to its free-living counterpart. Grey points indicate insignificant shift in selection, and red and blue points represent significant intensification and relaxation of selection, respectively.

**Table 4.** Summary of RELAX estimates for each lichen-forming taxon. The number of SOGs classified as intensified (K > 1), relaxed (K < 1), or not significant based on Benjamini–Hochberg adjusted p < 0.05; n indicates the total number of SOGs with valid estimates per taxa. Genes with |ln(K)| >10 were removed.

| Lichen-forming Taxa | n genes | n Intensified | n Relaxed | n Not Significant |
| --- | --- | --- | --- | --- |
| *Coccomyxa viridis* | 934 | 5 | 14 | 915 |
| *Symbiochloris reticulata* | 931 | 36 | 16 | 879 |
| *Trebouxia sp. C0010* | 941 | 105 | 7 | 829 |
| *Asterochloris erici* | 946 | 12 | 19 | 915 |

GO enrichment analysis of genes classified as significantly intensified or relaxed in each lineage recovered only one significant term. “Intracellular organelle lumen” was enriched among genes under intensified selection in *Symbiochloris reticulata* (p = 0.0057). No other GO terms were significantly enriched in any lineage across the Biological Process, Molecular Function, or Cellular Component gene ontologies.

To determine if any genes experience uniform intensification or relaxation of selection across lichen-forming algae, SOGs with K estimates in all four taxa (n= 906) were examined. Of these, 190 genes showed variable patterns across lineages (e.g., K > 1 in one lineage, K < 1 in another, not significant in others), and 715 genes were consistently non-significant across all four lineages. A single gene showed mostly intensified selection (K > 1 in three of four lineages; *Coccomyxa viridis*, *Trebouxia sp.* C0010 and *Asterochloris erici*). Two domains in this protein were identified with HHpred (Zimmermann et al., 2018) as a serine/threonine protein kinase domain (Pfam PF00069) and a class II aminoacyl-tRNA-synthetase-like domain (Pfam PF00152) (both probability = 100%). No genes showed uniformly intensified or relaxed selection across all four lineages.

**Alt text:** Volcano plots of shift in selection intensity. Most genes are grey and nonsignificant. *Trebouxia* and *Symbiochloris* show a larger red cluster than blue cluster, indicating more genes under intensified rather than relaxed selection. *Coccomyxa* and *Asterochloris* show few significant genes.

## Discussion

### Lichen symbiosis does not produce a uniform genomic syndrome

Lichen-forming algae have heterogeneous signatures of molecular evolution across all four metrics examined. Selection on synonymous codon usage is reduced in some lichen-forming taxa and intensified in others, and the intensity of selection on protein coding sequence varies both in magnitude and direction across lineages. This heterogeneity mirrors the pattern, or lack thereof, documented in other eukaryotic symbioses. Some show signs of genome reduction (termite-fungal symbionts and microsporidia; Conlon et al., 2021; Keeling & Slamovits, 2004) and/or elevated rates of molecular evolution (microsporidia; Haag et al., 2020), while others have slower rates of molecular evolution (lichen-forming fungi; Ametrano et al., 2022) or show sequence and structural divergence with hypothesized demographic drivers (coral-dinoflagellates; González-Pech et al., 2019, 2021). Lichen-forming algae fit squarely within this ambiguity.

### Lineage-specific ecology predicts variation in genomic response

The observed heterogeneity across signatures of molecular evolution likely reflects differences in the demographic and ecological contexts of each of the lichen-forming algae presented here. Different fungal partners likely create distinct physiological environments (e.g. variation in thallus architecture, water relations, secondary chemistry, or optical properties) which impose unique selective pressures on their associated algae. Differences in transmission mode, whether vertical, horizontal or mixed modes, likely modulate bottleneck severity (Bright & Bulgheresi, 2010; Mira & Moran, 2002; Peek et al., 1998). These lichen-forming algae also vary in their free-living capacity. The specific *Coccomyxa* examined here is considered “lichen-associated” and assumed to also exist free-living (Tagirdzhanova et al., 2023), and lichen-forming *Symbiochloris* and *Trebouxia* are found free-living on bark and soil, whereas *Asterochloris* appears more obligately lichenized (Sanders & Masumoto, 2021; Veselá et al., 2024).

The presence of free-living populations does not necessarily buffer against Ne reduction, because Ne determination is dependent on gene flow and recombination between and within free-living and lichenized populations. Recombination can cause Ne to vary across the genome, with areas of high recombination exhibiting population dynamics of higher Ne relative to areas of low recombination (Charlesworth, 2009). Although sexual reproduction is generally thought to be suppressed when lichenized (Ahmadjian, 1993; Friedl & Büdel, 2008; Hill, 1989, 2009; Sanders, 2023), lichen-forming algae likely are capable of sex when free-living. In *Trebouxia*, functional meiosis genes have been found (Gazquez et al., 2024; Tagirdzhanova et al., 2026); however, low admixture in *T. decolorans* populations suggests recombination could be rare (Wyczanska et al., 2023). These observations notwithstanding, many gaps remain in our understanding of the life-history of lichen-forming algae, particularly pertaining to their free-living phases. Examining Ne and its proxies can inform hypotheses about whether and how symbiosis affects the demography of lichen-forming algae.

### ω heterogeneity argues against uniform Ne constraint

Across lichen-forming algae, ω shifted in opposing directions. Two pairs exhibited elevated ω in lichen-forming lineages (*Coccomyxa* and *Asterochloris*, indicative of reduced efficacy of purifying selection or relaxed selective constraint, i.e. relaxed selection) and two showed reduced ω (*Symbiochloris* and *Trebouxia*, indicative of intensified selection) (Fig. 2B, Table 2). dN/dS is an established proxy for Ne across a diversity of lineages (Galtier, 2016; James et al., 2016; Popadin et al., 2007), and elevated substitution rates are a hallmark of demographic constraint in many endosymbiotic bacteria (McCutcheon & Moran, 2012; Moran, 1996; Woolfit & Bromham, 2003). However, because the direction of ω varies across lichen-forming algae, Ne reduction is not likely to be the primary factor influencing the molecular evolution of lichen-forming algae.

The decomposition of ω into its component parts gives pause to the apparent signal of intensified selection in *Trebouxia* and *Symbiochloris*. In *Trebouxia*, dN increased, consistent with reduced efficacy of selection, but dS increased even more, meaning the lower ω in *Trebouxia* is potentially reflective of CUB erosion rather than intensified protein-level selection. In *Symbiochloris*, both dN and dS shifted in directions that reduce ω relative to its free-living counterpart. CUB erosion alone cannot fully account for the reduced ratio; the concurrent decrease in dN is consistent with more effective purifying selection contributing to the reduced ω.

### Codon bias erosion is lineage-specific and may explain dS heterogeneity

Similar to rates of molecular evolution, signatures of CUB varied across taxon comparisons. *Symbiochloris* and *Trebouxia* showed increased ENC’ indicative of reduced CUB and *Coccomyxa* showed decreased ENC’, indicative of increased CUB, though the magnitude of this shift raises doubts about its biological significance. *Asterochloris* showed no significant change (Fig. 3A, Table 3). In contrast to this heterogeneity, a uniform reduction in GC3 in all lichen-forming taxa was found; however, an examination of the cause of this uniform reduction again suggests lineage-specific differences. Because selection on synonymous codon usage acts primarily at third codon positions, comparing shifts in GC3 to shifts in GC12 can distinguish reductions in the efficacy of selection at synonymous sites from genome-wide changes in mutational spectrum (Galtier et al., 2018; Sueoka, 1988). In *Symbiochloris*, GC12 did not significantly differ between lifestyles. In *Asterochloris*, GC12 shifted by <1% (ΔGC12 = −0.0023), which is negligible relative to the shift in GC3. The pattern in *Symbiochloris* and *Asterochloris* is thus consistent with reduced GC3 being driven by the reduced efficacy of selection on synonymous codon usage (Bulmer, 1991; Sharp et al., 2010). In *Coccomyxa* and *Trebouxia*, GC12 was significantly lower in lichen-forming taxa, indicating a genome-wide shift in the underlying mutational spectrum contributes to the observed GC3 decrease (Fig 3B, Table 3). In *Trebouxia*, both GC12 and GC3 decreased, indicating a contribution from a change in directional mutational pressure genome wide (Sueoka, 1988); however, the shift at GC12 was small relative to that at GC3, suggesting that reduced efficacy of selection on synonymous sites also contributes to the observed GC3 decrease. In *Coccomyxa*, the GC12 shift was ∼42% of the GC3 shift, but the small absolute magnitudes of both shifts make it difficult to attribute the GC3 decrease to a change in either the underlying mutational spectrum or selection efficacy at synonymous sites.

The reduced CUB observed in lichen-forming *Symbiochloris* and *Trebouxia* provides a potential explanation for why dS values are higher than in their free-living counterparts, challenging the standard assumption that synonymous substitutions are neutral. As originally described, dS reflects the neutral mutation rate (Kimura, 1977); however, in reality, dS includes both truly neutral sites and sites under weak selection (Chamary et al., 2006; Hershberg & Petrov, 2008; Plotkin & Kudla, 2011; Rahman et al., 2021). When Ne is large enough, selection is effective even for small selection coefficients, such as those associated with differences in codon usage (Bulmer, 1991; Sharp et al., 2010). When Ne drops below a critical threshold (|Nes| << 1; Kimura, 1983), synonymous sites that were under weak selection can become effectively neutral, causing dS to rise towards the neutral mutation rate (Rahman et al., 2021).

This raises the possibility that lichen-forming *Symbiochloris* and *Trebouxia* have experienced demographic constraints sufficient to erode selection on codon usage, while stronger purifying selection on nonsynonymous sites remains effective. This interpretation assumes more biased codon usage is under selection. In microbial symbioses such as lichens where organisms are in very close proximity, reduced CUB could be adaptive if tRNAs are exchanged between species; however, evidence for cross-species tRNA exchange is limited (e.g. Van Leuven et al., 2019).

The covariation of ΔENC’ and ΔdS is consistent with the interpretation that dS heterogeneity is driven in part by lineage-specific differences in the efficacy of selection on synonymous codon usage. *Trebouxia*, which had the largest ΔENC’, also had the largest ΔdS. *Symbiochloris*, which had a small but significant ΔENC’, had the second largest ΔdS. The two lineages with negative ΔENC’, *Coccomyxa* and *Asterochloris*, are the same lineages with a negative and insignificant ΔdS, respectively. Independent of Ne, shifts in dS can be influenced by other lineage-specific differences such as generation time, DNA repair, UV or desiccation stress in lichen thalli, all of which also affect mutation accumulation (Lutzoni & Pagel, 1997). GC-biased gene conversion (gBGC) could also contribute to the observed shift in GC3, which is a process where G/C alleles are favored over A/T alleles during recombination (Duret & Galtier, 2009; Mugal et al., 2015). Because sexual reproduction is generally thought to be suppressed when lichenized (Ahmadjian, 1993; Friedl & Büdel, 2008; Hill, 1989, 2009; Sanders, 2023), lichen-forming algae may experience reduced gBGC, causing GC3 to be lower independent of drift (Galtier et al., 2018). However, because GC12 is stable in some lineages (*Symbiochloris, Asterochloris*) but not others (*Coccomyxa, Trebouxia*), if gBGC is responsible, it likely contributes variably across lineages rather than acting as a uniform driver of GC3 reduction.

### Protein-level selection shows gene-specific and lineage-specific responses

RELAX estimates a selection intensity parameter (K) that estimates whether the distribution of ω across sites has shifted toward neutrality (K < 1) or away from it (K > 1). K cannot distinguish between reductions in Ne, which weaken the efficacy of selection at all sites, and changes in selection coefficients at specific loci (relaxed selection) (Wertheim et al., 2015). Of genes that were identified to have a significant shift in the strength of selection, *Symbiochloris* and *Trebouxia* had a significant excess of genes under intensified selection, which is consistent with the reduced ω observed in these lineages (Fig. 4). This suggests stronger purifying or positive selection; however, because K is estimated from the distribution of ω values, the elevated dS observed in these taxa complicates this interpretation. This apparent intensification could be reflective of CUB variation. Because no lineage showed relaxed selection genome-wide, the significant shifts detected by RELAX are more likely to reflect changes in selection coefficients at specific loci in response to symbiosis rather than a genome-wide reduction in the efficacy of selection driven by decreased Ne. With no genes consistently under intensified or relaxed selection, selective responses to symbiosis are likely not uniform but vary in lineage-specific ways.

## Limitations

There are a few limitations that arise from the phylogenetic framework used in this study that warrant caution in the interpretation of the presented molecular signatures. Identified SOGs across evolutionarily distant Trebouxiophyte taxa are inherently enriched for conserved genes, which are more likely to be essential or under stronger purifying selection than the genome as a whole. In these genes, most new mutations are likely to be strongly deleterious, such that Ne would need to be correspondingly low for most new mutations to fluctuate with the dynamics of drift. Taking this into account, this study provides a conservative test of signatures associated with the reduced efficacy of selection, and the absence of a uniform signal is not necessarily evidence that Ne reduction has not occurred.

The paired design was chosen to support a more sensitive test of how lichen-forming algal genomes compare to free-living genomes, as broad lifestyle comparisons across taxa are likely to mask (as this paper showed) variation in the genomic manifestations of lichenization. This required each lichen-forming taxon to have a sufficiently close, adequately sequenced free-living relative, which was available for few lichen-forming Trebouxiophyte alga, and absent for *Trebouxia* and *Asterochloris*, which both lack a free-living congener. While there are many sequenced free-living Trebouxiophyte algae, most are too phylogenetically distant from a lichen-forming counterpart to form an informative pair.

An additional important assumption in the interpretation of these results is that the free-living reference represents a demographic and selective baseline unaffected by lichenization. Current ancestral state reconstructions predict lichenization as the ancestral state in most of the Trebouxiophyceae (Puginier et al., 2024), raising the possibility that some “free-living” lineages retain genomic signatures of ancestral symbiosis. It is also possible that free-living algae experience demographic constraints unrelated to symbiosis. Ascribing results to lichenization therefore depends on the assumption that the unique life histories of free-living algae do not systematically bias the comparisons, and the results should be interpreted cautiously, particularly in light of their heterogeneity across free-living/lichen-forming pairs.

## Conclusion

The lack of a single genomic trajectory uncovered in lichen-forming algae (and other eukaryotic symbioses) likely results from a combination of key differences between many bacterial and eukaryotic symbioses. The most pronounced cases of reduced efficacy of selection are documented in endosymbiotic bacteria that exhibit strict vertical transmission, intracellular residence, and the absence of a free-living phase, which together result in extreme and sustained reductions in Ne (Bennett & Moran, 2015; Moran, 1996; Moran et al., 2008; Wernegreen, 2002). Most eukaryotic symbioses, including lichens, lack this combination of features, making it unsurprising that they do not show the same uniform genomic signatures. Beyond differences in the degree of demographic constraint, eukaryotic genomes may qualitatively differ from prokaryotic genomes in ways that affect how the reduced efficacy of selection manifests. Prokaryotic genomes possess a mutational bias towards deletion (Kuo et al., 2009; Kuo & Ochman, 2009), and when Ne drops, purifying selection cannot effectively oppose this bias, giving rise to endosymbiotic bacteria being some of the smallest genomes yet recorded (Moran & Bennett, 2014).

Eukaryotic genomes, by contrast, exhibit biases toward expansion through transposable element proliferation and segmental duplication (Lynch, 2007a; Lynch et al., 2011; Lynch & Conery, 2003; but see Lewin & Eyre-Walker, 2026; Marino et al., 2025; Weinstein & Roy, 2026), and possess fundamentally different genome architectures (linear chromosomes, variable ploidy) that may mediate the consequences of reduced Ne in ways that are not yet fully understood. Together, these differences in genome architecture, mutational bias, and the extremity of Ne reduction suggest that the “resident genome syndrome” (Andersson & Kurland, 1998) documented in many (but not all) bacterial endosymbionts cannot be straightforwardly extended to eukaryotic symbioses.

It is important to emphasize that the interpretation of these patterns across the molecular evolution landscape of lichen-forming algae is itself resting on the hypothesis that lichen symbiosis reduces Ne through transmission bottlenecks and/or reduced gene flow, with heterogeneity arising from variation in the severity of these constraints and lineage-specific shifts in the selective environment of each symbiosis. It is also in theory possible that algae residing within lichen thalli might actually have larger populations, could persist longer, and lead to larger Ne, as compared to ephemeral or patchy free-living algae (though this likely varies across different taxonomic groups and environments). Without direct Ne estimates from population data, the relative contributions of demographic constraint and adaptive evolution to these patterns remain unresolved. Lichen-forming algae, which vary along a continuum of free-living capacity, transmission mode, and potentially even symbiotic dependence, are an exciting system to explore how symbiosis affects the interacting influences of demography and selection on symbiotic genome evolution.

## Methods

### Genome Selection

Four lichen-forming/free-living comparisons were used: two within-genus comparisons (*Coccomyxa viridis* vs. C*occomyxa subellipsoidea*; *Symbiochloris reticulata* vs. *Symbiochloris irregularis*) and two cross-genus comparisons (*Trebouxia sp.* C0010 and *Asterochloris erici*, both compared to the free-living *Myrmecia bisecta*). *Trebouxia* and *Asterochloris* are both compared to *Myrmecia* because no free-living species in these genera are known. Within-genus comparisons provide the most direct evidence for lifestyle-associated evolutionary rate shifts. Cross-genus comparisons involve greater phylogenetic distance and longer independent evolutionary histories, which can introduce lineage-specific effects unrelated to lifestyle. Because *Myrmecia bisecta* serves as the free-living comparator for both *Trebouxia* and *Asterochloris*, these two comparisons are not fully independent; however, the contrasting results between these pairs (opposite directions of dN/dS and dS shifts) suggests differences in divergence time do not account for the heterogeneity in synonymous substitution rates between lichen-forming and free-living algae (see Results, Supplementary Fig. 3)

### Genome Annotation

Lichen-forming and free-living algal genomes were acquired from NCBI or JGI (Table 1). Genomes were softmasked for repetitive regions using RepeatMasker v4.1.1 with the Dfam 3.3 repeat library (Smit et al., 2013-2015). Softmasked genomes were annotated using the BRAKER3 pipeline, which uses the gene prediction tools GeneMark-EX and AUGUSTUS to perform an automated, semi-supervised gene annotation (Brůna et al., 2020, 2021; Buchfink et al., 2015; Gabriel et al., 2021; Gotoh et al., 2014; Iwata & Gotoh, 2012; Lomsadze et al., 2005; Stanke et al., 2006, 2008). BRAKER3 was run using an algal genome file and the Viridiplantae OrthoDB database with the --softmasking and – min_contig=1000 flags to train GeneMark-EX and AUGUSTUS. The AUGUSTUS flag -- genemodel=complete was used to restrict the output annotation to only complete protein sequences. Annotation completeness was assessed with BUSCO v5.4.3 (Manni et al., 2021) in protein mode on BRAKER3 predicted protein files, against the chlorophyta_odb10 lineage dataset. Complete BUSCOs ranged from 86.4% to 94.1% across taxa (Supplementary Table 1).

### Phylogenomic Reconstruction

OrthoFinder v2.5.4 was used with default parameters to identify single-copy orthologous genes (SOGs) shared across all taxa (Emms & Kelly, 2019). For each SOG, predicted protein sequences from the BRAKER3 annotations were aligned with MUSCLE v3.8.31 (Edgar, 2004), and codon-based nucleotide alignments were generated by backtranslating each protein alignment against its corresponding coding sequences with PAL2NAL v14 (Suyama et al., 2006). Alignments were then trimmed to remove poorly aligned regions using TrimAl with the –gappyout algorithm (Capella-Gutiérrez et al., 2009). Gene trees were subsequently constructed in IQ-TREE v.2.1.2, using the substitution model LG+F+G4, which was selected after first running model testing (-m TEST) (Minh et al., 2020) on a representative SOG alignment. The inferred gene-tree topologies were used by PAML and RELAX, which reestimate branch lengths and selection parameters using their own codon substitution models. A majority rule consensus tree with mean branch lengths was constructed with the SumTrees program within the DendroPy Python package (v.4.0.0) (Sukumaran & Holder, 2010). Node values represent the proportion of gene trees containing each partition.

### Functional Annotation

Functional annotation of the identified protein coding regions (n=951) was performed with InterProScan v5.23-62.0 against the Pfam, PANTHER, Gene3D, SUPERFAMILY and CDD databases (Jones et al., 2014). InterPro domain assignments and associated Gene Ontology (GO) terms were parsed from the InterProScan TSV output and used for downstream GO enrichment analyses.

### dN/dS Analysis

To estimate dN/dS ratios (ω) for each SOG across algal taxa, the codeml program in the PAML package was used (Yang, 2007). Three models were tested: (i) a null model where a single ω was estimated across all branches, (ii) a two-ratio model where one ω value was estimated for lichen-forming branches (foreground) and one ω value was estimated for free-living branches (background), and (iii) a free-ratio model where a separate ω was estimated for each branch. For each model, codeml was run with the parameters CodonFreq = 2, clock = 0, NSsites = 0, fix_omega = 0, omega = 0.4, and cleandata = 0. The specific gene tree generated with IQ-TREE was used for each sequence analyzed. Branch specific dN, dS, ω, and log-likelihood values were extracted for each SOG. All downstream analyses were performed in R v4.6.0.

Model fit was evaluated for each SOG with likelihood ratio tests comparing the log-likelihood of the null model to the two-ratio and free-ratio models. The two-ratio model outperformed the null model in 23.1% of SOGs, whereas the free-ratio model outperformed the null model in 62.6% of SOGs; the free-ratio model was therefore used for all downstream analyses (see Supplementary Fig. 4 for a comparison of pooled lifestyle effects under the two models). Before further statistical analysis, SOGs that were saturated with or had too few substitutions to reliably estimate ω were removed (dS < 0.01, dS > 3, dN/dS > 10). The upper dS bound follows Ametrano et al. (2022) and is more stringent than the ω based filtering used in many comparable analyses (Harrison et al., 2024; Rubin & Moreau, 2016). The lower bound (dS > 0.01; Toll-Riera et al., 2011) excludes branches with too few synonymous substitutions, for which ω estimates are unreliable. A sensitivity analysis (Supplementary Fig. 2) confirmed that estimated marginal mean differences in dS and ω between taxon pairs were stable across dS filtering thresholds of 2, 3, and 5. Differences in dN were stable except in *Symbiochloris* and *Trebouxia*, where the lifestyle effect reached significance at the dS < 3 threshold. For genera with multiple available genome assemblies (*Asterochloris*, *Trebouxia*), one representative genome was chosen at random for the main analyses. Including multiple closely related sequences from the same genus produced phylogenies with highly uneven branch lengths, which drove dS estimates on the short branches toward zero yielding unreliable ω estimates. A single representative per genus was also more consistent with the paired comparative design. The effect of genome choice on estimated marginal mean differences between taxon pairs was assessed in a sensitivity analysis using each available assembly separately (Supplementary Fig. 1).

For each response (dN, dS, ω), a linear mixed effects model was fit as follows using the lme4 package (Bates et al., 2015): response ∼ condition × taxon pair + (1|SOG). SOG was modeled as a random intercept to account for nonindependence among branches within orthologs. Degrees of freedom reported are Satterthwaite approximations generated with the lmerTest package (Kuznetsova et al., 2017). Residual diagnostics indicated departure from normality in the upper tail, consistent with the right skewed nature of these molecular evolution metrics. Log transformation did not improve residual diagnostics, but because inferences from linear mixed models are robust to moderate departures from normality (Schielzeth et al., 2020), dN, dS, and ω were analyzed on the original scale. Pairwise contrasts of estimated marginal means within each taxon pair were obtained from this model using the emmeans package (Lenth & Piaskowski 2026) with Bonferroni correction (Dunn, 1961) across the four pairs. Pooled lifestyle effects across pairs were obtained as marginal means from the same model. The same modelling approach was applied to ENC’, GC12 and GC3.

### Codon Use Bias Analysis

CUB was quantified from codon-based alignments for each SOG. For each alignment, the Effective Number of Codons (ENC’) was calculated using the codonbias Python package (Diament 2022). ENC’ is more robust to cross species comparisons as it controls for background variation in GC content (Novembre, 2002). GC content was computed separately for the first, second, and third codon positions (GC1, GC2, GC3). Linear mixed effects models using all SOGS (n=951) were parameterized as described for dN/dS analyses in R v4.6.0.

### RELAX Analysis

Before performing the RELAX analysis, each gene tree was labeled for four separate analyses, each with the focal lichen-forming alga labeled as the test branch and the respective free-living alga as the reference branch. The input phylogeny contained all taxa to maintain the same topology across analyses. RELAX v. 4.5 (Wertheim et al., 2015) was then run separately on each SOG within each comparison (951 SOGs × 4 comparisons = 3,804 analyses) using the -srv Yes flag to account for synonymous rate variation. The relaxation parameter K and its associated p-value were extracted from each analysis. Final gene numbers for each lineage differed due to RELAX failing to converge for some genes in each lineage (Table 4). All downstream analyses were performed in R v4.6.0.

Before comparing K values across taxa, p-values were Benjamini-Hochberg adjusted (Benjamini & Hochberg, 1995) and any SOG with |ln(K)| > 10 was flagged as an extreme K estimate and removed. The number of genes for which selection was intensified (K >1; p < 0.05), relaxed (K <1; p < 0.05) and not significant (p > 0.05) was tabulated. To test whether any taxa had a significant bias towards relaxation or intensification of selection, a two-sided exact binomial test was performed against the null expectation of equal proportion of genes under relaxation and intensification of selection. P-values were Bonferroni adjusted across the four lineages. Any gene under uniform relaxation or intensification of selection in all four lichen-forming taxa was labeled “consistently relaxed/intensified”, any gene under relaxation or intensification of selection in three of four taxa was labeled “mostly relaxed/intensified”, any gene under relaxation or intensification of selection in two of four taxa was labeled “variable”. Functional annotation of the gene under intensified selection was performed with HHpred (MPI Bioinformatics Toolkit; Zimmermann et al., 2018). The query sequence was searched against the PDB70, Pfam-A, SMART, and SCOPe70 profile HMM databases using the toolkit’s default HHsearch parameters.

### Gene Ontology (GO) Enrichment Analysis

Using GO terms inferred with InterProScan a GO enrichment analysis was performed using the topGO R package v.3.23 (Alexa & Rahnenfuhrer 2026). For each of the four lichen-forming taxa, two enrichment tests were performed across each of the three GO ontologies (Biological Process, Molecular Function, Cellular Component). One test was performed with genes classified as significantly relaxed and a second testing genes classified as significantly intensified. For each analysis, the background gene set was constructed from all SOGs for which RELAX successfully estimated K in that taxon and for which at least one GO annotation was identified. Enrichment was tested using the “weight01” algorithm with Fisher’s exact test. GO terms with fewer than 5 genes were removed from the analysis. P-values were Benjamini-Hochberg adjusted to correct for multiple hypothesis testing (Benjamini & Hochberg, 1995). Results were considered significant at an adjusted p-value < 0.05. All GO terms and ontology classifications were retrieved from the GO.db R package (Carlson et al., 2025).

## Data Availability

This study used publicly available genome assemblies downloaded from NCBI and JGI; accession numbers and assembly codes are provided in Table 1. Custom scripts for sequence extraction, alignment, and downstream statistical analyses, along with intermediate data files (per-gene dN/dS, ENC′, GC, and RELAX estimates), are available at github.com/anonymouslichen/Algae-Comparative-Genomics and will be archived at Zenodo upon acceptance. No new sequence data were generated in this study.

## Acknowledgements

A sincere thank you to my advisor and committee members, Daniel Stanton, R. Ford Denison, Ruth Shaw and Amelia Lindsey whose conversations and comments greatly improved the ideas presented here. Thank you to Matthew Nelson, with whom conversations during the early conception of this study influenced the trajectory of this work and to members of the Stanton Lab for their comments on previous versions of this manuscript.

## References

Ahmadjian, V. (1993). The lichen symbiosis. John Wiley.

Álvarez-Carretero, S., Kapli, P., & Yang, Z. (2023). Beginner’s Guide on the Use of PAML to Detect Positive Selection. Molecular Biology and Evolution, 40(4), msad041. 10.1093/molbev/msad041

Ametrano, C. G., Lumbsch, H. T., Di Stefano, I., Sangvichien, E., Muggia, L., & Grewe, F. (2022). Should we hail the Red King? Evolutionary consequences of a mutualistic lifestyle in genomes of lichenized ascomycetes. Ecology and Evolution, 12(1), e8471. 10.1002/ece3.8471

Andersson, S. G. E., & Kurland, C. G. (1998). Reductive evolution of resident genomes. Trends in Microbiology, 6(7), 263–268. 10.1016/S0966-842X(98)01312-2

Armaleo, D., Müller, O., Lutzoni, F., Andrésson, Ó. S., Blanc, G., Bode, H. B., Collart, F. R., Dal Grande, F., Dietrich, F., Grigoriev, I. V., Joneson, S., Kuo, A., Larsen, P. E., Logsdon, J. M., Lopez, D., Martin, F., May, S. P., McDonald, T. R., Merchant, S. S., … Xavier, B. B. (2019). The lichen symbiosis re-viewed through the genomes of Cladonia grayi and its algal partner Asterochloris glomerata. BMC Genomics, 20(1), 605. 10.1186/s12864-019-5629-x

Bates, D., Mächler, M., Bolker, B., & Walker, S. (2015). Fitting Linear Mixed-Effects Models Using **lme4**. Journal of Statistical Software, 67(1). 10.18637/jss.v067.i01

Benjamini, Y., & Hochberg, Y. (1995). Controlling the False Discovery Rate: A Practical and Powerful Approach to Multiple Testing. Journal of the Royal Statistical Society Series B: Statistical Methodology, 57(1), 289–300. 10.1111/j.2517-6161.1995.tb02031.x

Bennett, G. M., & Moran, N. A. (2015). Heritable symbiosis: The advantages and perils of an evolutionary rabbit hole. Proceedings of the National Academy of Sciences, 112(33), 10169– 10176. 10.1073/pnas.1421388112

Bright, M., & Bulgheresi, S. (2010). A complex journey: Transmission of microbial symbionts. Nature Reviews Microbiology, 8(3), 218–230. 10.1038/nrmicro2262

Bromham, L. (2009). Why do species vary in their rate of molecular evolution? Biology Letters, 5(3), 401–404. 10.1098/rsbl.2009.0136

Brůna, T., Hoff, K. J., Lomsadze, A., Stanke, M., & Borodovsky, M. (2021). BRAKER2: Automatic eukaryotic genome annotation with GeneMark-EP+ and AUGUSTUS supported by a protein database. NAR Genomics and Bioinformatics, 3(1), lqaa108. 10.1093/nargab/lqaa108

Brůna, T., Lomsadze, A., & Borodovsky, M. (2020). GeneMark-EP+: Eukaryotic gene prediction with self-training in the space of genes and proteins. NAR Genomics and Bioinformatics, 2(2), lqaa026. 10.1093/nargab/lqaa026

Buchfink, B., Xie, C., & Huson, D. H. (2015). Fast and sensitive protein alignment using DIAMOND. Nature Methods, 12(1), 59–60. 10.1038/nmeth.3176

Bulmer, M. (1991). The selection-mutation-drift theory of synonymous codon usage. Genetics, 129(3), 897–907. 10.1093/genetics/129.3.897

Capella-Gutiérrez, S., Silla-Martínez, J. M., & Gabaldón, T. (2009). trimAl: A tool for automated alignment trimming in large-scale phylogenetic analyses. Bioinformatics, 25(15), 1972–1973. 10.1093/bioinformatics/btp348

Chamary, J. V., Parmley, J. L., & Hurst, L. D. (2006). Hearing silence: Non-neutral evolution at synonymous sites in mammals. Nature Reviews Genetics, 7(2), 98–108. 10.1038/nrg1770

Charlesworth, B. (2009). Effective population size and patterns of molecular evolution and variation. Nature Reviews Genetics, 10(3), 195–205. 10.1038/nrg2526

Conlon, B. H., Gostinčar, C., Fricke, J., Kreuzenbeck, N. B., Daniel, J.-M., Schlosser, M. S. L., Peereboom, N., Aanen, D. K., De Beer, Z. W., Beemelmanns, C., Gunde-Cimerman, N., & Poulsen, M. (2021). Genome reduction and relaxed selection is associated with the transition to symbiosis in the basidiomycete genus Podaxis. iScience, 24(6), 102680. 10.1016/j.isci.2021.102680

Douglas, A. E. (2016). How multi-partner endosymbioses function. Nature Reviews Microbiology, 14(12), 731–743. 10.1038/nrmicro.2016.151

Dunn, O. J. (1961). Multiple Comparisons among Means. Journal of the American Statistical Association, 56(293), 52–64. 10.1080/01621459.1961.10482090

Duret, L., & Galtier, N. (2009). Biased Gene Conversion and the Evolution of Mammalian Genomic Landscapes. Annual Review of Genomics and Human Genetics, 10(1), 285–311. 10.1146/annurev-genom-082908-150001

Edgar, R. C. (2004). MUSCLE: Multiple sequence alignment with high accuracy and high throughput. Nucleic Acids Research, 32(5), 1792–1797. 10.1093/nar/gkh340

Emms, D. M., & Kelly, S. (2019). OrthoFinder: Phylogenetic orthology inference for comparative genomics. Genome Biology, 20(1), 238. 10.1186/s13059-019-1832-y

Friedl, T., & Büdel, B. (2008). Photobionts. In T. H. Nash (Ed.), Lichen Biology (2nd ed., pp. 9–26). Cambridge University Press. 10.1017/CBO9780511790478.003

Gabriel, L., Hoff, K. J., Brůna, T., Borodovsky, M., & Stanke, M. (2021). TSEBRA: transcript selector for BRAKER. BMC Bioinformatics, 22(1), 566. 10.1186/s12859-021-04482-0

Galtier, N. (2016). Adaptive Protein Evolution in Animals and the Effective Population Size Hypothesis. PLOS Genetics, 12(1), e1005774. 10.1371/journal.pgen.1005774

Galtier, N., Roux, C., Rousselle, M., Romiguier, J., Figuet, E., Glemin, S., Bierne, N., & Duret, L. (2018). Codon Usage Bias in Animals: Disentangling the Effects of Natural Selection, Effective Population Size, and GC-Biased Gene Conversion. Molecular Biology and Evolution, 35(5), 1092–1103. 10.1093/molbev/msy015

Gazquez, A., Bordenave, C. D., Montero-Pau, J., Pérez-Rodrigo, M., Marco, F., Martínez-Alberola, F., Muggia, L., Barreno, E., & Carrasco, P. (2024). From spores to gametes: A sexual life cycle in a symbiotic *Trebouxia* microalga. Algal Research, 84, 103744. 10.1016/j.algal.2024.103744

González-Pech, R. A., Bhattacharya, D., Ragan, M. A., & Chan, C. X. (2019). Genome Evolution of Coral Reef Symbionts as Intracellular Residents. Trends in Ecology & Evolution, 34(9), 799–806. 10.1016/j.tree.2019.04.010

González-Pech, R. A., Stephens, T. G., Chen, Y., Mohamed, A. R., Cheng, Y., Shah, S., Dougan, K. E., Fortuin, M. D. A., Lagorce, R., Burt, D. W., Bhattacharya, D., Ragan, M. A., & Chan, C. X. (2021). Comparison of 15 dinoflagellate genomes reveals extensive sequence and structural divergence in family Symbiodiniaceae and genus Symbiodinium. BMC Biology, 19(1), 73. 10.1186/s12915-021-00994-6

Gotoh, O., Morita, M., & Nelson, D. R. (2014). Assessment and refinement of eukaryotic gene structure prediction with gene-structure-aware multiple protein sequence alignment. BMC Bioinformatics, 15(1), 189. 10.1186/1471-2105-15-189

Haag, K. L., Pombert, J.-F., Sun, Y., De Albuquerque, N. R. M., Batliner, B., Fields, P., Lopes, T. F., & Ebert, D. (2020). Microsporidia with Vertical Transmission Were Likely Shaped by Nonadaptive Processes. Genome Biology and Evolution, 12(1), 3599–3614. 10.1093/gbe/evz270

Harrison, T. L., Stinchcombe, J. R., & Frederickson, M. E. (2024). Elevated Rates of Molecular Evolution Genome-wide in Mutualist Legumes and Rhizobia. Molecular Biology and Evolution, 41(12), msae245. 10.1093/molbev/msae245

Hershberg, R., & Petrov, D. A. (2008). Selection on Codon Bias. Annual Review of Genetics, 42(1), 287–299. 10.1146/annurev.genet.42.110807.091442

Hill, D. J. (1989). The control of the cell cycle in microbial symbionts. New Phytologist, 112(2), 175–184. 10.1111/j.1469-8137.1989.tb02372.x

Hill, D. J. (2009). Asymmetric Co-evolution in the Lichen Symbiosis Caused by a Limited Capacity for Adaptation in the Photobiont. The Botanical Review, 75(3), 326–338. 10.1007/s12229-009-9028-x

Husnik, F., & McCutcheon, J. P. (2018). Functional horizontal gene transfer from bacteria to eukaryotes. Nature Reviews Microbiology, 16(2), 67–79. 10.1038/nrmicro.2017.137

Iwata, H., & Gotoh, O. (2012). Benchmarking spliced alignment programs including Spaln2, an extended version of Spaln that incorporates additional species-specific features. Nucleic Acids Research, 40(20), e161–e161. 10.1093/nar/gks708

James, J. E., Piganeau, G., & Eyre-Walker, A. (2016). The rate of adaptive evolution in animal mitochondria. Molecular Ecology, 25(1), 67–78. 10.1111/mec.13475

Jones, P., Binns, D., Chang, H.-Y., Fraser, M., Li, W., McAnulla, C., McWilliam, H., Maslen, J., Mitchell, A., Nuka, G., Pesseat, S., Quinn, A. F., Sangrador-Vegas, A., Scheremetjew, M., Yong, S.-Y., Lopez, R., & Hunter, S. (2014). InterProScan 5: Genome-scale protein function classification. Bioinformatics, 30(9), 1236–1240. 10.1093/bioinformatics/btu031

Keeling, P. J., & Palmer, J. D. (2008). Horizontal gene transfer in eukaryotic evolution. Nature Reviews Genetics, 9(8), 605–618. 10.1038/nrg2386

Keeling, P. J., & Slamovits, C. H. (2004). Simplicity and Complexity of Microsporidian Genomes. Eukaryotic Cell, 3(6), 1363–1369. 10.1128/EC.3.6.1363-1369.2004

Kimura, M. (1977). Preponderance of synonymous changes as evidence for the neutral theory of molecular evolution. Nature, 267(5608), 275–276. 10.1038/267275a0

Kimura, M. (1983). The Neutral Theory of Molecular Evolution. Cambridge University Press. Cambridge Core. 10.1017/CBO9780511623486

Koch, N. M., Lendemer, J. C., Manzitto-Tripp, E. A., McCain, C., & Stanton, D. E. (2023). Carbon-concentrating mechanisms are a key trait in lichen ecology and distribution. Ecology, 104(5), e4011. 10.1002/ecy.4011

Kuo, C.-H., Moran, N. A., & Ochman, H. (2009). The consequences of genetic drift for bacterial genome complexity. Genome Research, 19(8), 1450–1454. 10.1101/gr.091785.109

Kuo, C.-H., & Ochman, H. (2009). Deletional Bias across the Three Domains of Life. Genome Biology and Evolution, 1, 145–152. 10.1093/gbe/evp016

Kuznetsova, A., Brockhoff, P. B., & Christensen, R. H. B. (2017). **lmerTest** Package: Tests in Linear Mixed Effects Models. Journal of Statistical Software, 82(13). 10.18637/jss.v082.i13

Lewin, L., & Eyre-Walker, A. (2026). A Comparative Analysis of Long-Term Effective Population Sizes Across Eukaryotes. Molecular Ecology, 35(4), e70265. 10.1111/mec.70265

Lomsadze, A., Ter-Hovhannisyan, V., Chernoff, Y. O., & Borodovsky, M. (2005). Gene identification in novel eukaryotic genomes by self-training algorithm. Nucleic Acids Research, 33(20), 6494– 6506. 10.1093/nar/gki937

Lutzoni, F., & Pagel, M. (1997). Accelerated evolution as a consequence of transitions to mutualism. Proceedings of the National Academy of Sciences, 94(21), 11422–11427. 10.1073/pnas.94.21.11422

Lynch, M. (2007a). The frailty of adaptive hypotheses for the origins of organismal complexity. Proceedings of the National Academy of Sciences, 104(suppl_1), 8597–8604. 10.1073/pnas.0702207104

Lynch, M. (2007b). The origins of genome architecture. Sinauer Associates.

Lynch, M., Bobay, L.-M., Catania, F., Gout, J.-F., & Rho, M. (2011). The Repatterning of Eukaryotic Genomes by Random Genetic Drift. Annual Review of Genomics and Human Genetics, 12(1), 347–366. 10.1146/annurev-genom-082410-101412

Lynch, M., & Conery, J. S. (2003). The Origins of Genome Complexity. Science, 302(5649), 1401–1404. 10.1126/science.1089370

Manni, M., Berkeley, M. R., Seppey, M., & Zdobnov, E. M. (2021). BUSCO: Assessing Genomic Data Quality and Beyond. Current Protocols, 1(12), e323. 10.1002/cpz1.323

Marino, A., Debaecker, G., Fiston-Lavier, A.-S., Haudry, A., & Nabholz, B. (2025). Effective population size does not explain long-term variation in genome size and transposable element content in animals. 10.7554/eLife.100574.2

McCutcheon, J. P., & Moran, N. A. (2012). Extreme genome reduction in symbiotic bacteria. Nature Reviews Microbiology, 10(1), 13–26. 10.1038/nrmicro2670

Minh, B. Q., Schmidt, H. A., Chernomor, O., Schrempf, D., Woodhams, M. D., Von Haeseler, A., & Lanfear, R. (2020). IQ-TREE 2: New Models and Efficient Methods for Phylogenetic Inference in the Genomic Era. Molecular Biology and Evolution, 37(5), 1530–1534. 10.1093/molbev/msaa015

Mira, A., & Moran, N. A. (2002). Estimating Population Size and Transmission Bottlenecks in Maternally Transmitted Endosymbiotic Bacteria. Microbial Ecology, 44(2), 137–143. 10.1007/s00248-002-0012-9

Moran, N. A. (1996). Accelerated evolution and Muller’s rachet in endosymbiotic bacteria. Proceedings of the National Academy of Sciences, 93(7), 2873–2878. 10.1073/pnas.93.7.2873

Moran, N. A. (2007). Symbiosis as an adaptive process and source of phenotypic complexity. Proceedings of the National Academy of Sciences, 104(suppl_1), 8627–8633. 10.1073/pnas.0611659104

Moran, N. A., & Bennett, G. M. (2014). The Tiniest Tiny Genomes. Annual Review of Microbiology, 68(1), 195–215. 10.1146/annurev-micro-091213-112901

Moran, N. A., McCutcheon, J. P., & Nakabachi, A. (2008). Genomics and Evolution of Heritable Bacterial Symbionts. Annual Review of Genetics, 42(1), 165–190. 10.1146/annurev.genet.41.110306.130119

Mugal, C. F., Weber, C. C., & Ellegren, H. (2015). GC-biased gene conversion links the recombination landscape and demography to genomic base composition. BioEssays, 37(12), 1317–1326. 10.1002/bies.201500058

Nelsen, M. P., Leavitt, S. D., Heller, K., Muggia, L., & Lumbsch, H. T. (2021). Macroecological diversification and convergence in a clade of keystone symbionts. FEMS Microbiology Ecology, 97(6), fiab072. 10.1093/femsec/fiab072

Nelsen, M. P., Leavitt, S. D., Heller, K., Muggia, L., & Lumbsch, H. T. (2022). Contrasting Patterns of Climatic Niche Divergence in Trebouxia—A Clade of Lichen-Forming Algae. Frontiers in Microbiology, 13. https://www.frontiersin.org/articles/10.3389/fmicb.2022.791546

Novembre, J. A. (2002). Accounting for Background Nucleotide Composition When Measuring Codon Usage Bias. Molecular Biology and Evolution, 19(8), 1390–1394. 10.1093/oxfordjournals.molbev.a004201

Ohta, T. (1992). The Nearly Neutral Theory of Molecular Evolution. Annual Review of Ecology and Systematics, 23, 263–286.

Peek, A. S., Vrijenhoek, R. C., & Gaut, B. S. (1998). Accelerated Evolutionary Rate in Sulfur-Oxidizing Endosymbiotic Bacteria Associated with the Mode of Symbiont Transmission. Molecular Biology and Evolution, 15(11), 1514–1523. 10.1093/oxfordjournals.molbev.a025879

Plotkin, J. B., & Kudla, G. (2011). Synonymous but not the same: The causes and consequences of codon bias. Nature Reviews Genetics, 12(1), 32–42. 10.1038/nrg2899

Popadin, K., Polishchuk, L. V., Mamirova, L., Knorre, D., & Gunbin, K. (2007). Accumulation of slightly deleterious mutations in mitochondrial protein-coding genes of large versus small mammals. Proceedings of the National Academy of Sciences, 104(33), 13390–13395. 10.1073/pnas.0701256104

Poquita-Du, R. C., Otte, J., Calchera, A., & Schmitt, I. (2024). Genome-Wide Comparisons Reveal Extensive Divergence Within the Lichen Photobiont Genus, *Trebouxia*. Genome Biology and Evolution, 16(10), evae219. 10.1093/gbe/evae219

Puginier, C., Libourel, C., Otte, J., Skaloud, P., Haon, M., Grisel, S., Petersen, M., Berrin, J.-G., Delaux, P.-M., Dal Grande, F., & Keller, J. (2024). Phylogenomics reveals the evolutionary origins of lichenization in chlorophyte algae. Nature Communications, 15(1), 4452. 10.1038/s41467-024-48787-z

Rahman, S., Kosakovsky Pond, S. L., Webb, A., & Hey, J. (2021). Weak selection on synonymous codons substantially inflates *dN/dS* estimates in bacteria. Proceedings of the National Academy of Sciences, 118(20), e2023575118. 10.1073/pnas.2023575118

Rubin, B. E. R., & Moreau, C. S. (2016). Comparative genomics reveals convergent rates of evolution in ant–plant mutualisms. Nature Communications, 7(1), 12679. 10.1038/ncomms12679

Russell, S. L., Corbett-Detig, R. B., & Cavanaugh, C. M. (2017). Mixed transmission modes and dynamic genome evolution in an obligate animal–bacterial symbiosis. The ISME Journal, 11(6), 1359– 1371. 10.1038/ismej.2017.10

Russell, S. L., Pepper-Tunick, E., Svedberg, J., Byrne, A., Ruelas Castillo, J., Vollmers, C., Beinart, R. A., & Corbett-Detig, R. (2020). Horizontal transmission and recombination maintain forever young bacterial symbiont genomes. PLOS Genetics, 16(8), e1008935. 10.1371/journal.pgen.1008935

Sagan, L. (1967). On the origin of mitosing cells. Journal of Theoretical Biology, 14(3), 225,IN1-274,IN6.

Sanders, W. B. (2023). Is lichen symbiont mutualism a myth? BioScience, 73(9), 623–634. 10.1093/biosci/biad073

Sanders, W. B., & Masumoto, H. (2021). Lichen algae: The photosynthetic partners in lichen symbioses. The Lichenologist, 53(5), 347–393. 10.1017/S0024282921000335

Schielzeth, H., Dingemanse, N. J., Nakagawa, S., Westneat, D. F., Allegue, H., Teplitsky, C., Réale, D., Dochtermann, N. A., Garamszegi, L. Z., & Araya-Ajoy, Y. G. (2020). Robustness of linear mixed-effects models to violations of distributional assumptions. Methods in Ecology and Evolution, 11(9), 1141–1152. 10.1111/2041-210X.13434

Sharp, P. M., Emery, L. R., & Zeng, K. (2010). Forces that influence the evolution of codon bias. Philosophical Transactions of the Royal Society B: Biological Sciences, 365(1544), 1203–1212. 10.1098/rstb.2009.0305

Stanke, M., Diekhans, M., Baertsch, R., & Haussler, D. (2008). Using native and syntenically mapped cDNA alignments to improve de novo gene finding. Bioinformatics, 24(5), 637–644. 10.1093/bioinformatics/btn013

Stanke, M., Schöffmann, O., Morgenstern, B., & Waack, S. (2006). Gene prediction in eukaryotes with a generalized hidden Markov model that uses hints from external sources. BMC Bioinformatics, 7(1), 62. 10.1186/1471-2105-7-62

Sudakaran, S., Kost, C., & Kaltenpoth, M. (2017). Symbiont Acquisition and Replacement as a Source of Ecological Innovation. Trends in Microbiology, 25(5), 375–390. 10.1016/j.tim.2017.02.014

Sueoka, N. (1988). Directional mutation pressure and neutral molecular evolution. Proceedings of the National Academy of Sciences, 85(8), 2653–2657. 10.1073/pnas.85.8.2653

Sukumaran, J., & Holder, M. T. (2010). DendroPy: A Python library for phylogenetic computing. Bioinformatics, 26(12), 1569–1571. 10.1093/bioinformatics/btq228

Suyama, M., Torrents, D., & Bork, P. (2006). PAL2NAL: Robust conversion of protein sequence alignments into the corresponding codon alignments. Nucleic Acids Research, 34(Web Server), W609–W612. 10.1093/nar/gkl315

Tagirdzhanova, G., Scharnagl, K., Sahu, N., Yan, X., Bucknell, A., Bentham, A. R., Jégousse, C., Ament-Velásquez, S. L., Onuț-Brännström, I., Johannesson, H., MacLean, D., & Talbot, N. J. (2025). Complexity of the lichen symbiosis revealed by metagenome and transcriptome analysis of Xanthoria parietina. Current Biology, 35(4), 799–817.e5. 10.1016/j.cub.2024.12.041

Tagirdzhanova, G., Scharnagl, K., Yan, X., & Talbot, N. J. (2023). Genomic analysis of Coccomyxa viridis, a common low-abundance alga associated with lichen symbioses. Scientific Reports, 13(1), Article 1. 10.1038/s41598-023-48637-w

Toll-Riera, M., Laurie, S., & Alba, M. M. (2011). Lineage-Specific Variation in Intensity of Natural Selection in Mammals. Molecular Biology and Evolution, 28(1), 383–398. 10.1093/molbev/msq206

Van Leuven, J. T., Mao, M., Xing, D. D., Bennett, G. M., & McCutcheon, J. P. (2019). Cicada Endosymbionts Have tRNAs That Are Correctly Processed Despite Having Genomes That Do Not Encode All of the tRNA Processing Machinery. mBio, 10(3), e01950–18. 10.1128/mBio.01950-18

Veselá, V., Malavasi, V., & Škaloud, P. (2024). A synopsis of green-algal lichen symbionts with an emphasis on their free-living lifestyle. Phycologia, 63(3), 317–338. 10.1080/00318884.2024.2325329

Weinstein, B., & Roy, S. W. (2026). Controlling for life-history traits in vertebrates reveals that effective population size does not affect mutation rate or genome size. Proceedings of the National Academy of Sciences, 123(6), e2519649123. 10.1073/pnas.2519649123

Wernegreen, J. J. (2002). Genome evolution in bacterial endosymbionts of insects. Nature Reviews Genetics, 3(11), Article 11. 10.1038/nrg931

Wertheim, J. O., Murrell, B., Smith, M. D., Kosakovsky Pond, S. L., & Scheffler, K. (2015). RELAX: Detecting Relaxed Selection in a Phylogenetic Framework. Molecular Biology and Evolution, 32(3), 820–832. 10.1093/molbev/msu400

Woolfit, M., & Bromham, L. (2003). Increased Rates of Sequence Evolution in Endosymbiotic Bacteria and Fungi with Small Effective Population Sizes. Molecular Biology and Evolution, 20(9), 1545– 1555. 10.1093/molbev/msg167

Wyczanska, M., Wacker, K., Dyer, P. S., & Werth, S. (2023). Local-scale panmixia in the lichenized fungus Xanthoria parietina contrasts with substantial genetic structure in its Trebouxia photobionts. The Lichenologist, 1–11. 10.1017/S002428292300004X

Xiong, Q., Zheng, L., Zhang, Q., Li, T., Zheng, L., & Song, L. (2025). Comparative genomic insights into ecological adaptations and evolutionary dynamics of Trebouxiophyceae algae. BMC Genomics, 26(1), 764. 10.1186/s12864-025-11933-y

Yang, Z. (2007). PAML 4: Phylogenetic Analysis by Maximum Likelihood. Molecular Biology and Evolution, 24(8), 1586–1591. 10.1093/molbev/msm088

Zoller, S., & Lutzoni, F. (2003). Slow algae, fast fungi: Exceptionally high nucleotide substitution rate differences between lichenized fungi Omphalina and their symbiotic green algae Coccomyxa. Molecular Phylogenetics and Evolution, 29(3), 629–640. Red

