## Supplemental Figures & Table for "Lichen symbiosis does not impose a uniform genomic syndrome on algae"

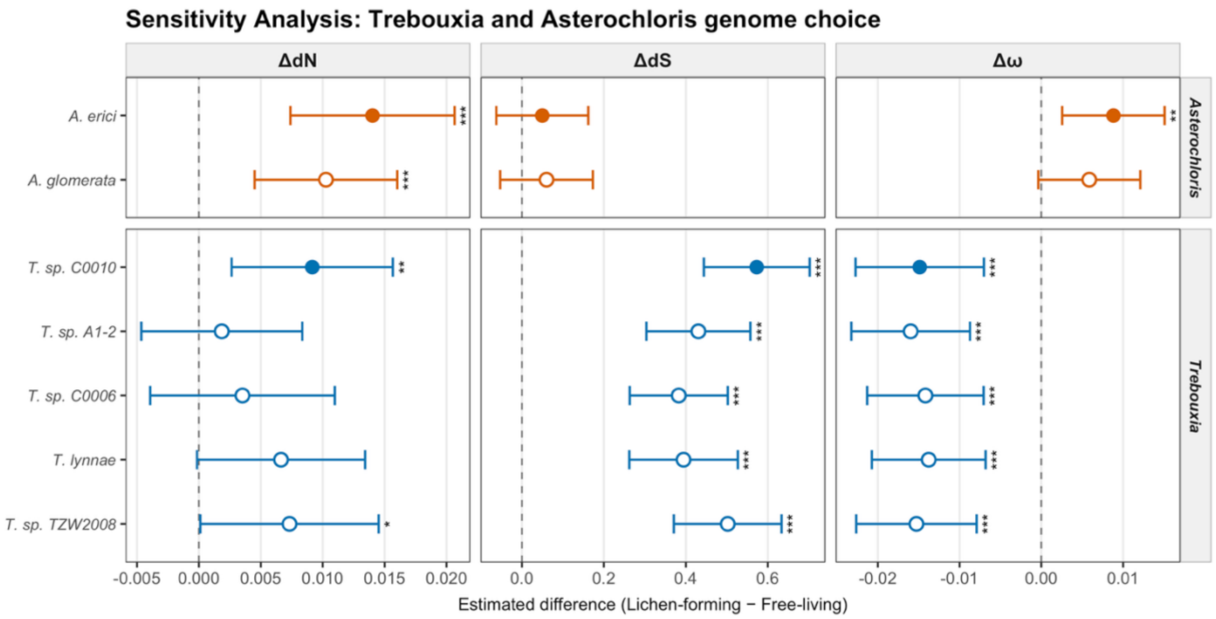


**Supplementary Figure 1.** Effect of genome choice on estimated differences of absolute and relative rates of molecular evolution for *Asterochloris* and *Trebouxia*, for which multiple genomes were available. Points indicate estimated marginal mean differences between each *Asterochloris* and *Trebouxia* genome and their shared free-living counterpart, *Myrmecia bisecta*, indicating the reported results are robust to bias introduced by genome choice. Bars represent the 95% CI around each estimate. Solid circle indicates the genomes used and estimates reported in main study findings. * = p<0.05, ** = p<0.01, and *** = p<0.001.

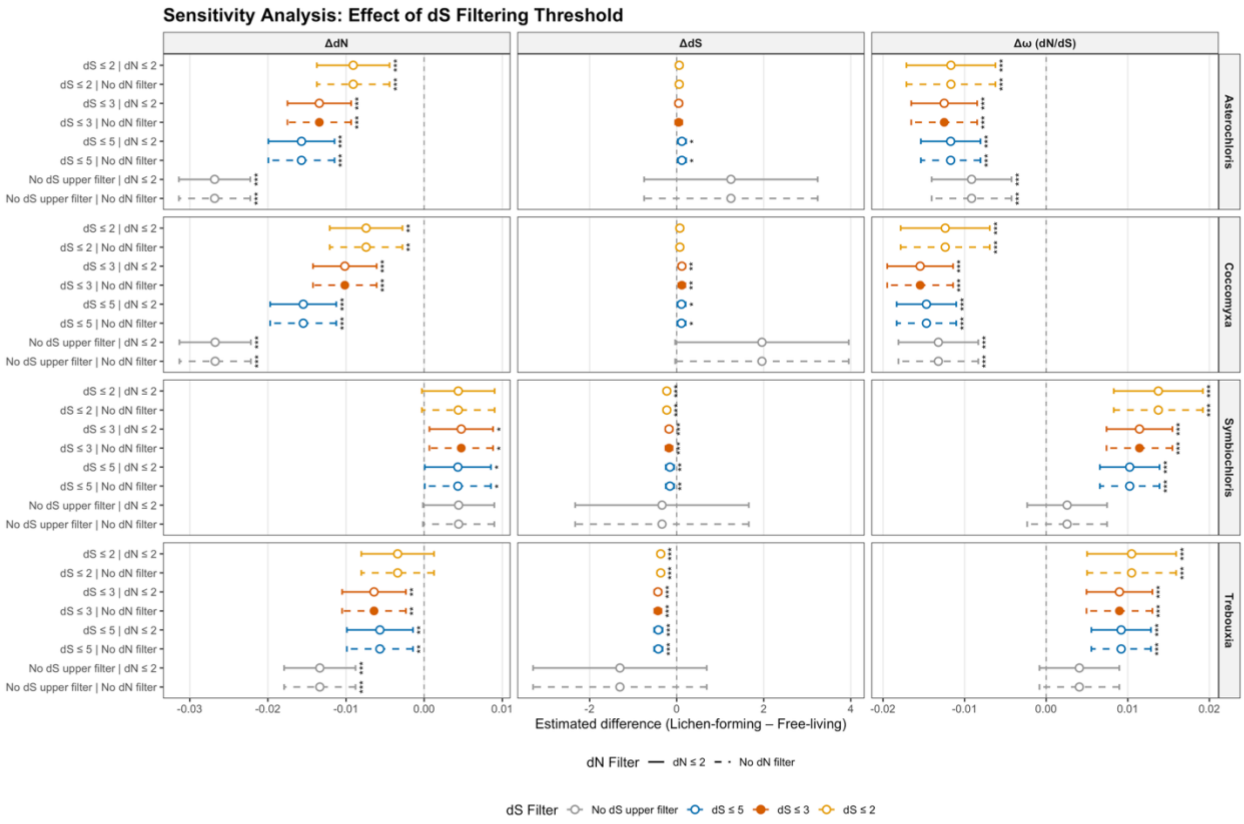


**Supplementary Figure 2.** Effect of dS filtering threshold on estimated differences on absolute and relative rates of molecular evolution. Points indicate estimated marginal mean differences between lichen-forming and free-living taxa. Bars represent the 95% CI around each estimate. For all sensitivity analyses include only dS > 0.01 and omega < 10 (standard in the literature). Colors indicate variation in upper dS filtering threshold (2, 3, 5, no upper bound). Solid lines indicate dN <= 2 (Toll-Riera et al. 2011), dashed line indicates no dN filter. Solid circle indicates the filtering threshold and estimates reported in main study findings.  * = p<0.05, ** =  p<0.01, and *** = p<0.001.

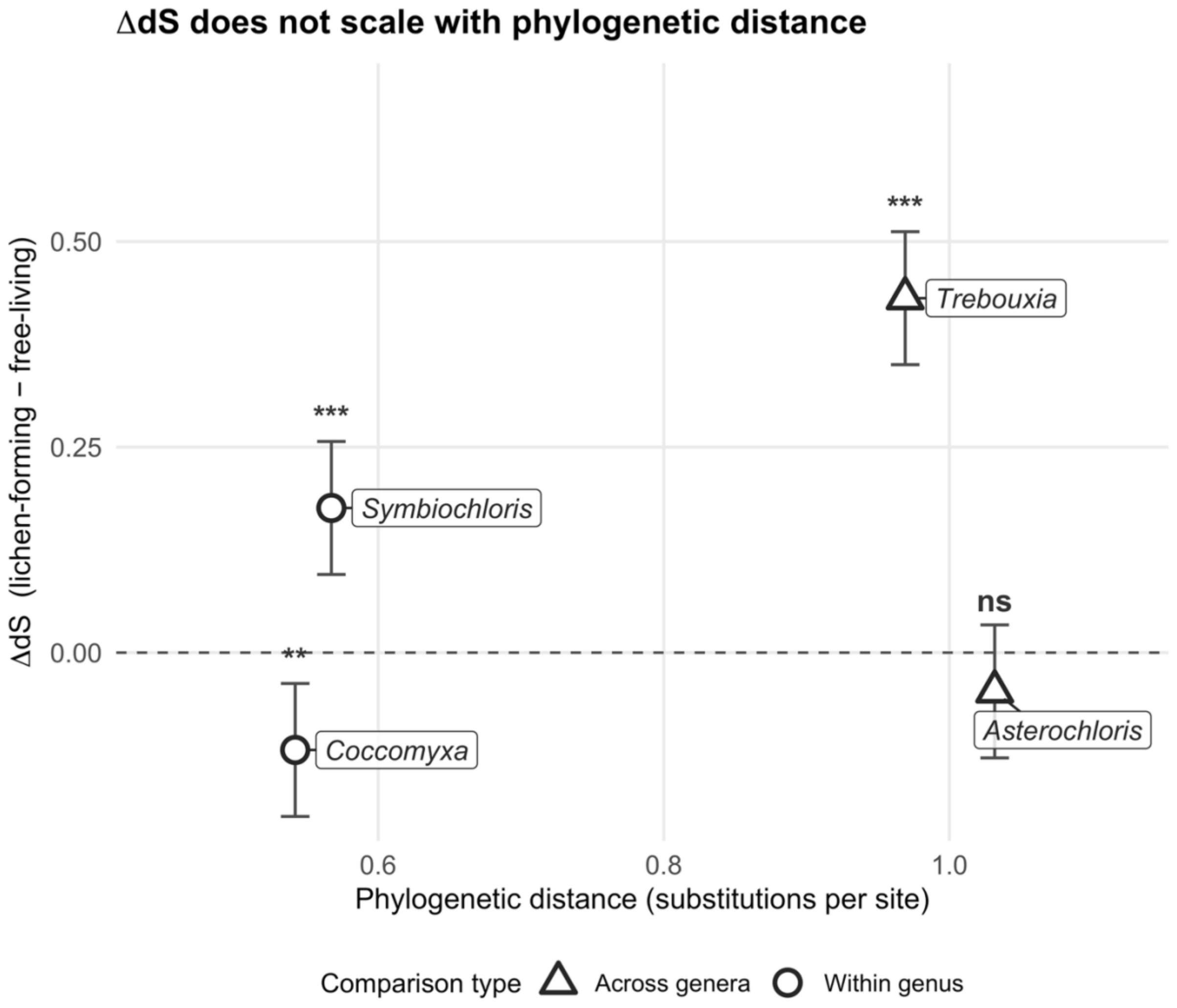


**Supplementary Figure 3.** Relationship between phylogenetic distance and ΔdS (lichen-forming - free-living) across taxon pairs. Phylogenetic distances are from the consensus species tree. ΔdS values and 95% confidence intervals are from pairwise contrasts of estimated marginal means from a linear mixed-effects model. Within-genus comparisons are indicated by circles and cross-genus comparisons are indicated by triangles. The two most phylogenetically distant pairs have opposite ΔdS, indicating that differences in divergence time do not drive heterogeneity in synonymous substitution rates between lichen-forming and free-living algae. Significance levels indicate whether ΔdS differs from zero for each pair (**$p < 0.01$, ***$p < 0.001$, ns = not significant).

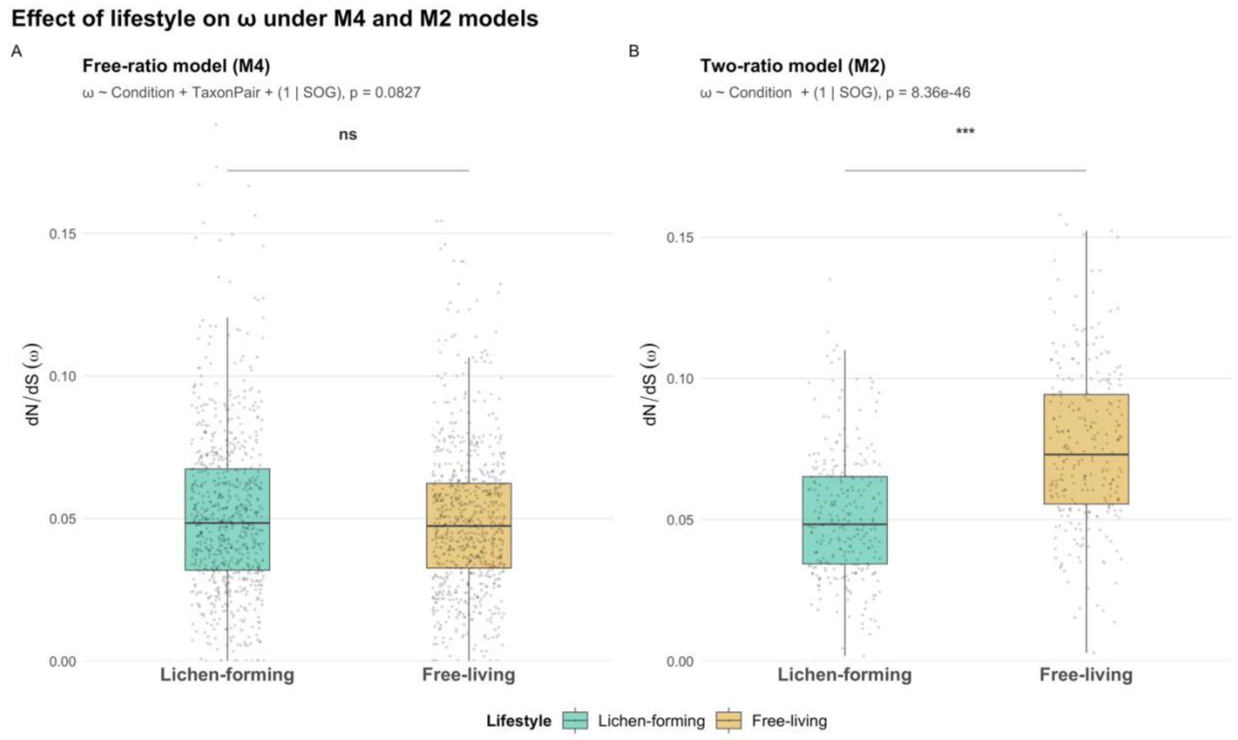


**Supplementary Figure 4.** Pooled comparisons of ω between lichen-forming and free-living algae under the (A) free-ratio model and (B) the two-ratio model. The two-ratio model estimates one ω for all lichen-forming branches and one ω for all free-living branches. This model yielded a significant effect of lifestyle (linear mixed-effects model with condition as fixed effect and SOG as a random intercept, $F_{1, 269} = 301.37$, $p < 0.0001$). This model outperformed the null model in only 23.1% of SOGs, compared to 62.6% for the free-ratio model. Under the free-ratio model, no significant overall effect of lifestyle was detected (linear mixed-effects model with condition and taxon pair as fixed effects and SOG as random intercepts, $F_{1,1291} = 3.016$, $p = 0.0827$). This discrepancy reflects the significant lifestyle × taxon pair interaction reported in the main results (linear mixed-effects model with condition × taxonpair fixed effect and SOG as a random intercept, $F_{3,1288} = 46.43$, $p < 0.0001$), indicating that opposing lineage-specific effects are masked when averaged across taxa.

| Species | Source | Assembly Code | Complete (%) | Single-copy (%) | Duplicated (%) | Fragmented (%) | Missing (%) | Total BUSCOs searched |
|---|---|---|---|---|---|---|---|---|
| *Coccomyxa viridis* | NCBI | GCA_964019345.2 | 94.1 | 82.1 | 12 | 0.1 | 5.8 | 1519 |
| *Coccomyxa subellipsoidea* | NCBI | GCF_000258705.1 | 91.9 | 73.3 | 18.6 | 0.5 | 7.6 | 1519 |
| *Symbiochloris reticulata* | JGI | 1016105 | 90 | 80.6 | 9.4 | 0.4 | 9.6 | 1519 |
| *Symbiochloris irregularis* | NCBI | GCA_040144405.1 | 89.3 | 78.1 | 11.2 | 0.5 | 10.2 | 1519 |
| *Trebouxia* sp. C0010 | NCBI | GCA_045269315.1 | 86.4 | 77 | 9.4 | 0.3 | 13.3 | 1519 |
| *Myrmecia bisecta* | NCBI | GCA_040144395.1 | 86.9 | 76 | 10.9 | 2.9 | 10.2 | 1519 |
| *Asterochloris erici* | NCBI | GCA_019693375.1 | 89.7 | 77.2 | 12.5 | 0.5 | 9.8 | 1519 |

**Supplementary Table 1.** BUSCO assessment (chlorophyta_odb10) of BRAKER3 predicted protein annotations for each genome.
